# HaploHide: A Data Hiding Framework for Privacy Enhanced Sharing of Personal Genetic Data

**DOI:** 10.1101/786517

**Authors:** Arif Harmanci, Xiaoqian Jiang, Degui Zhi

## Abstract

Personal genetic data is becoming a digital commodity as millions of individuals have direct access to and control of their genetic information. This information must be protected as it can be used for reidentification and potential discrimination of individuals and relatives. While there is a great incentive to share and use genetic information, there are limited number of practical approaches for protecting it when individuals would like to make use of their genomes in clinical and recreational settings. To enable privacy-enhanced usage of genomic data by individuals, we propose a crowd-blending-based framework where portions of the individual’s haplotype is “hidden” within a large sample of other haplotypes. The hiding framework is motivated by the existence of large-scale population panels that we utilize for generation of the crowd of haplotypes in which the individual’s haplotype is hidden. We demonstrate the usage of hiding in two different scenarios: Sharing of variant alleles on genes and sharing of GWAS variant alleles. We evaluate hiding framework by testing reidentification of hidden individuals using numerous measures of individual reidentification. In these settings, we discuss how effective hiding can be accomplished when the adversary does not have access to auxiliary identifying information. Compared to the existing approaches for protecting privacy, which require substantial changes in the computational infrastructure, e.g., homomorphic encryption, hiding-based framework does not incur any changes to the infrastructure. However, the processing must be performed for every sample in the crowd and therefore data processing cost will increase as the crowd size increases.

## Introduction

The advances in DNA sequencing technologies brought a substantial increase in the amount of personal genomic information. Starting with the initial population-wide genotyping projects such as The HapMap Consortium[1], The 1000 Genomes Project[2], Genomics England[3], The Cancer Genome Atlas (TCGA)[4], Trans-omics for precision medicine (TOPMed)[5], The Genotype-Tissue Expression (GTEx) Project[6], and the Precision Medicine Initiative[7], the number of available personal genomes has surpassed millions.

While these genetic data offer great opportunities to improve health and cure diseases, the sharing of the genetic variants brought up unprecedented challenges for privacy of individuals[8]–[12]. Many of these challenges are underpinned in the recreational use of DNA for ancestry tracing and in the clinics for genetic testing. In addition, as the clinical utility of genetic data is increasing, such as prenatal genetic testing[13] and disease risk prediction[14], the genetic testing is becoming more prevalent in many fronts. Many startup companies aim to capitalize on providing custom services such as ancestry search and disease risk estimation using individuals’ genetic information[15]. While individuals are sharing their genetic information, it is extremely challenging to keep the data perfectly safe because of the data breaches [16]. In addition, individuals may not trust the companies with their genetic data because of unclear and overly complex regulations around data sharing. In addition, these risks may extend to the relatives of the individuals [15] [16]. While individuals would like to make use of their genetic information, they may potentially face a conundrum between the eagerness to learn more about their genetics, health, and ancestry and the fear of discrimination. On another note, many individuals share their genetic information publicly on the online genetic genealogy portals. However, several recent studies have pointed out that genetic data is highly identifying and it can be used to link the individual’s information to unexpected databases[19], [20]. Moreover, recent advances in usage of DNA as incriminating forensic evidence to solve high-profile cases brings many ethical questions that may cause concerns for these individuals[21], [22].

The field of genomic privacy has grown substantially in the recent years. Numerous studies have shown that small number of variants can lead to re-identification attacks[8], [23], [24] and linking attacks[20], [25], [26]. One of the major frameworks that have been successfully applied to genomic data analysis is differential privacy[27], [28] (DP). In DP based approaches, data releasers design privacy-enabling data release mechanisms that generate aggregate statistics from data. To enable privacy, noise is added to the released data. While DP enables strong privacy-preserving guarantees (even when adversary has strong auxiliary information), the noise addition is a major drawback since the noise may corrupt certain data types (such as genotype data) to the extent that the data becomes unusable. In addition, DP based methods focus on statistical data release (e.g. mean, variance, aggregated statistics) and they are not immediately applicable for protecting the privacy of one individual (i.e., sample size of 1) who wants to share their genomic data. Currently the encryption-based approaches represent the most rigorous route to securely share personal genetic information[29]. There are, however, challenges about their practicality[30]. Methods such as homomorphic encryption[31] enable analyzing and processing encrypted data directly, however, there is a substantial computational cost and each algorithm must be re-implemented and optimized to efficiently analyze the encrypted data in the homomorphic manner. Moreover, the untrusted parties (i.e., a genetic testing company) may not be willing to convert all of their analysis systems to enable homomorphic encryption because of the involved storage, computational, and maintenance costs. Secondly, another major bottleneck is the know-how and technical competency required to maintain the encryption systems and encrypted databases. In this regard, key management must be performed securely to ensure the security and integrity of the encrypted databases. Another privacy-preserving data sharing formalism in this arena is secure multiparty computation (MPC) based systems[32]. These systems rely heavily on secret-sharing based approaches such that the data is divided into shares and each share is processed by independent data analyzers. MPC-based systems may also not be always practical since they rely on heavy communication and computational requirements on the parties[33]. In addition, non-collusion assumption between analyzers may not always hold and systems must be implemented in a problem specific manner. Due to the above-mentioned challenges, there is currently a lack of practical frameworks that will enable individuals to share their genetic information without involving high computational complexity with an untrusted third party and make use of it while not revealing their genetic information.

Here, we present HaploHide, a data hiding framework that can be used to enhance an individual’s privacy while sharing their genetic information with an untrusted third party. The basic idea behind HaploHide is to make use of existing publicly available genomic databases and hide an individual’s genetic information within a cohort of individuals, we term *metacohort*, such that an adversary cannot identify the individual’s genotypes among the samples, which we refer to as the *metasamples*. The metacohort represent a “crowd” and hiding procedure enables the individual to blend in this crowd. Within the hiding framework, the untrusted party, for example a genetic testing company, processes the genetic data for all the metasamples and returns the results back to the individual for all the metasamples, which include the result for the hidden individual. Since all the data that is exchanged is plain genomic data, the company does not have to perform any changes to their data analysis pipelines. At the same time, the individual (i.e. the customer) can simply discard the results for all the metasamples except her/his own. Thus, the data hiding approaches can be integrated seamlessly into the existing personal genomic data sharing approaches.

There have been several similar approaches to preserve individual privacy in other fields[34]. For example, in order to preserve location-privacy, a hiding based approach has been utilized to ensure that the untrusted parties cannot reliably identify an individual’s location[35], [36]. In this scenario, a user wants to use location-based services from an untrusted company to receive, for example, routing instructions between two locations that the user wants to navigate. To protect the location privacy of the user, authors propose “camouflaging” the user’s location among a crowd to ensure that the user’s location and destination cannot be reliably determined by an untrusted party. Another application was for protecting online privacy. This became particularly urgent when the regulation on internet service providers (ISPs) have been removed to collect information from individual user’s web browsing histories. This may create privacy concerns around ISPs sharing browsing histories with the 3^rd^ party companies such as advertisers. In the proposed approaches (such as “Internet Noise”[37] and “noiszy”[38]), web browser plugins are used to automatically visit random web sites. The random visits create a history of “noisy web browsing history” and the aim is to hide the user’s actual browsing history within the crowd of noisy web history. These approaches, however, are unlikely to provide meaningful privacy protection because the noisy web history can be relatively easily separated from the user’s actual web site visits. The main drawback stems from the fact that the untrusted 3^rd^ parties are really looking for identifying regularly visited sites so that they can reveal a user’s personal preferences. The noisy web sites do not help camouflage these regularly visited web sites since the noisy browsing history does not reflect a real individual’s browsing habits. Thus, a simple “attack” on this hiding scenario by measuring the frequency of web sites visits can easily filter out the noisy web history from the real web browsing habits of the user. This points out the fact that it is necessary to create the crowd in a realistic manner that simulates the real data.

In the genomic context, our approach employs a similar hiding-in-the-crowd strategy. HaploHide relies on the growing number of publicly available genomic databases such as The 1000 Genomes Project dataset[2], which represent a powerful knowledge-base for generating the crowd among which a real individual is hidden. We refer to this dataset as the metacohort generation panel. HaploHide makes use of the individuals in the generation panel to build a crowd that is indistinguishable from the real individual that is being hidden. As we discuss, the generation panel is of vital importance for effective hiding and the mismatches between genetic background of the hidden individual and the generation panel may decrease the effectiveness of hiding. This underscores the fact that the crowd must be generated properly to simulate a real cohort among which the real individual can hide. To properly simulate the metasamples in the crowd, our approach utilizes the generation panel directly by projecting the individual’s alleles on the panel or it uses the generation panel to estimate allele frequencies and sample variant alleles from the estimated frequencies. We demonstrate the use of these two approaches for sharing variant alleles on cancer-associated genes and the alleles of GWAS variants. One of the main components of our study is evaluating the effectiveness of hiding by considering attack models. For this, we test the practicality of hiding framework by evaluating a large number of statistics that the adversary can utilize to systematically re-identify the hidden individual. Overall, our results suggest that the hiding mechanism as a promising framework for privacy-enhanced sharing high dimensional genetic information.

## Results

We first summarize the data hiding scenario and the application of HaploHide algorithm. Then we present two applications of genomic data hiding framework.

### Data Hiding Scenario

Figure 1a illustrates the setup of the proposed hiding based framework. The individual, we refer to as Alice, would like to send her variant alleles to another party who will perform a computation and send results back to Alice. The other party is assumed to be a trusted but curious entity. In this example, we assume that the other party is the company that performs disease risk estimation using Alice’s variant alleles.

**Figure 1.**
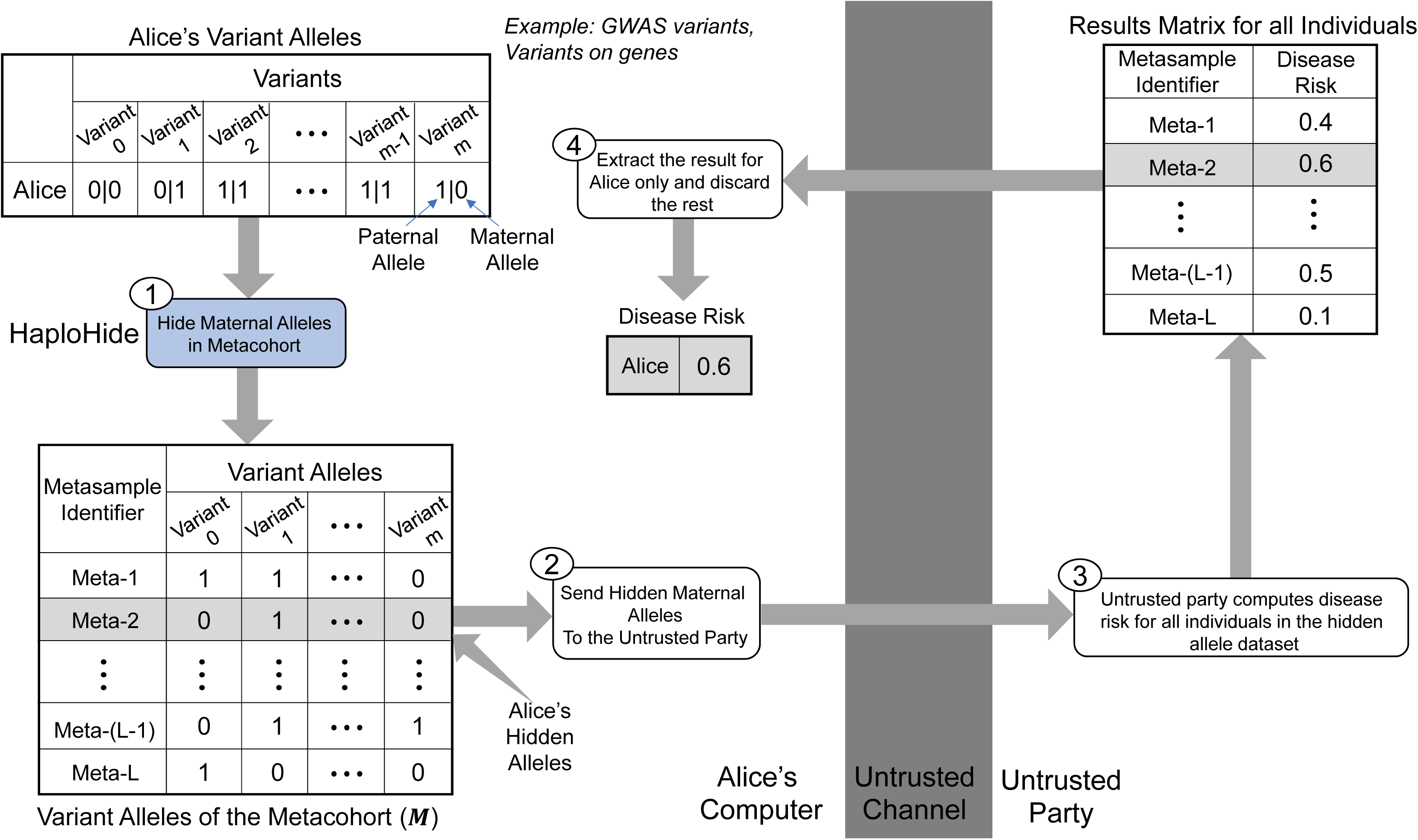
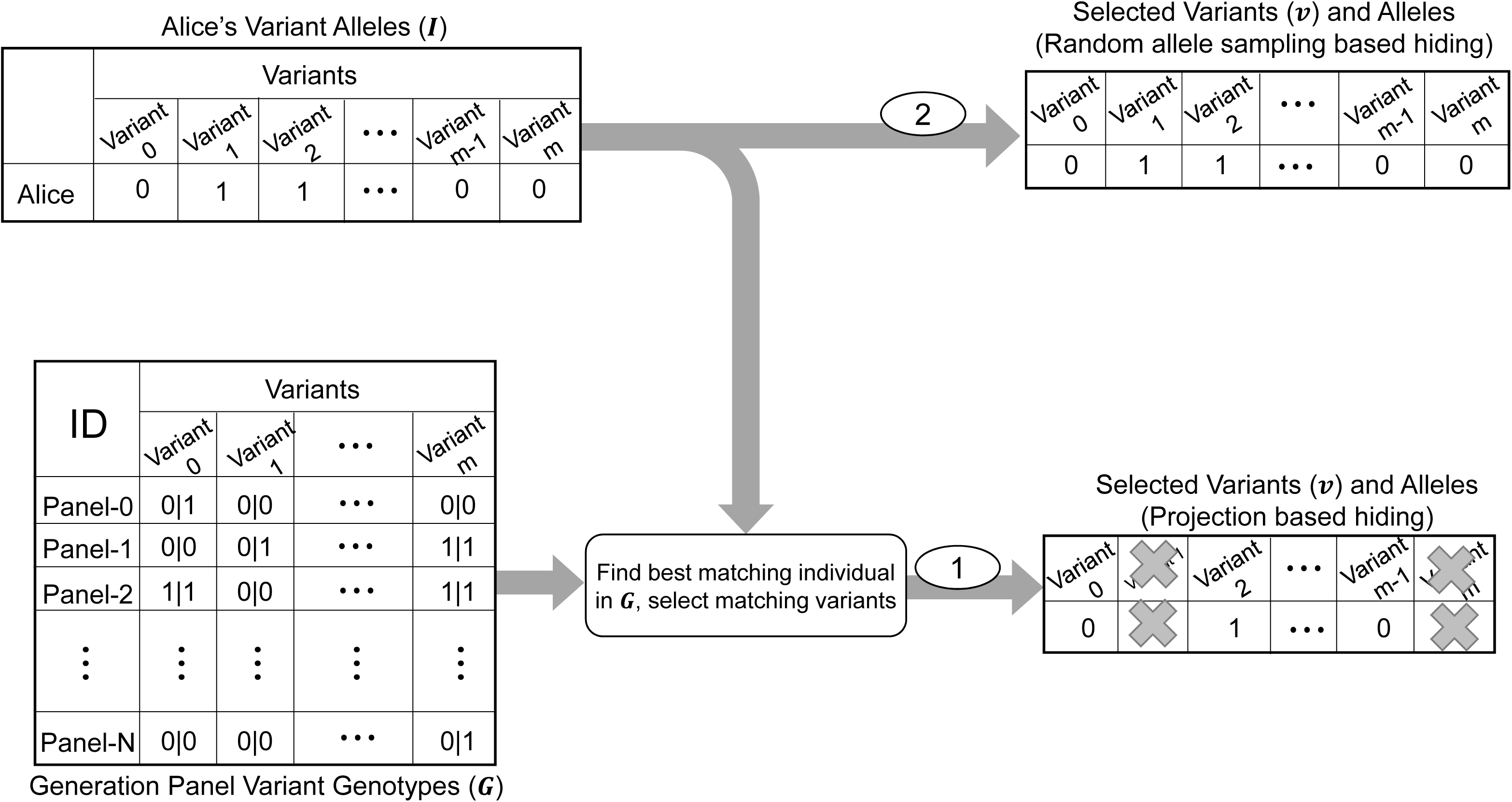
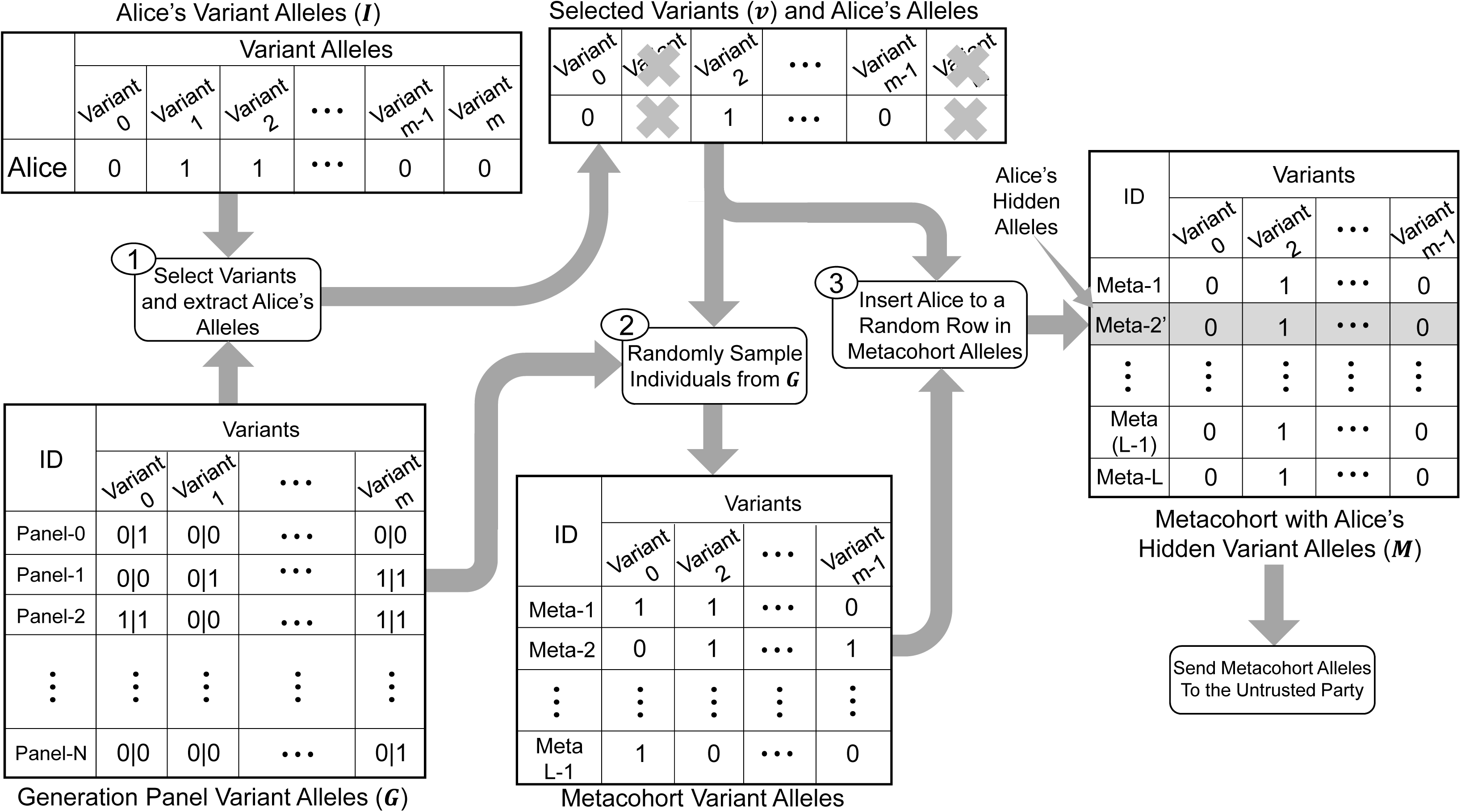
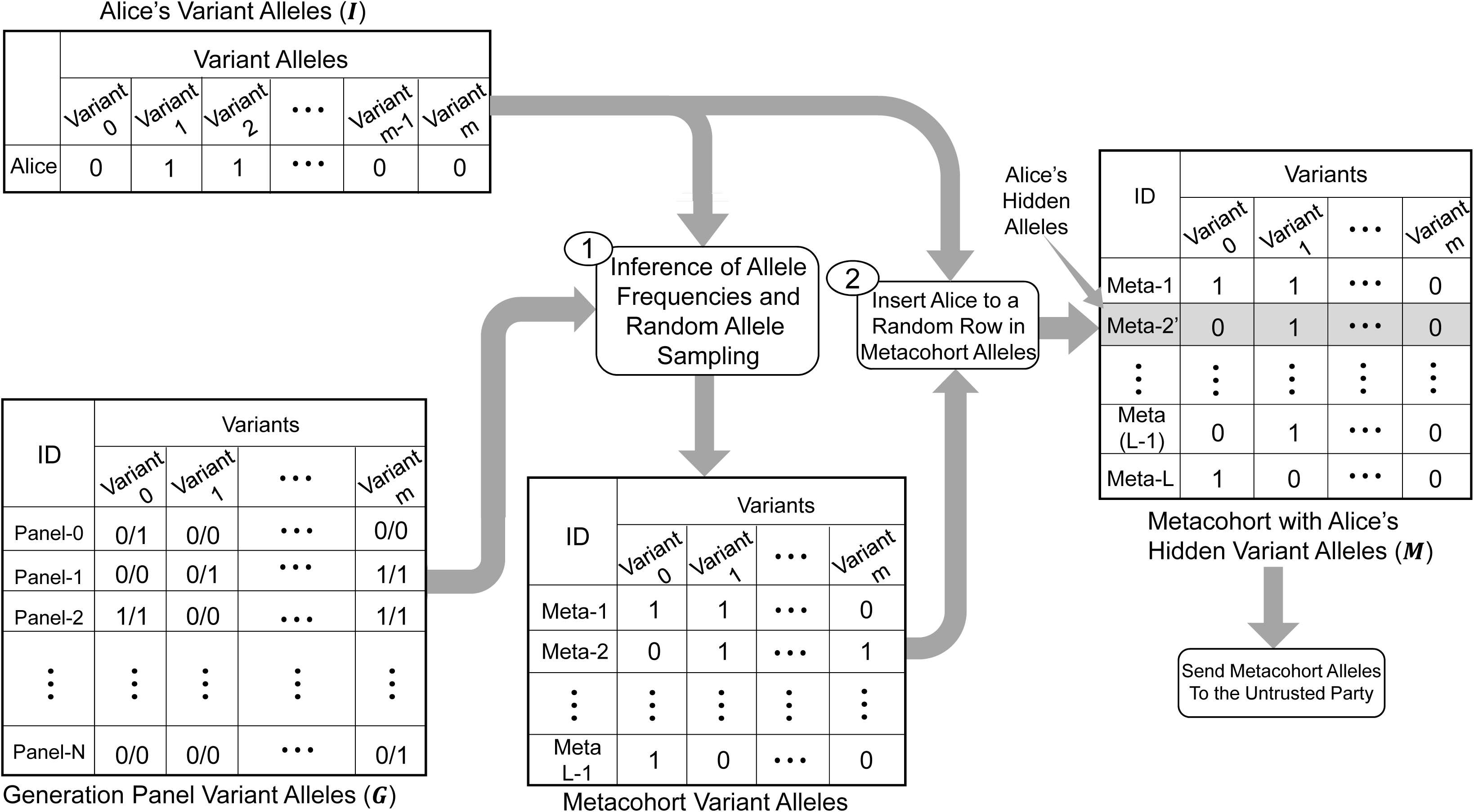
Illustration of the hiding framework in a disease risk computation scenario. ***a)*** There are *m* variant alleles (denoted by ***I***) that Alice would like to send to the untrusted party. Alice and the untrusted party are connected by an untrusted channel. In step 1, the Alice’s variants are extracted and are hidden using HaploHide. In the example, Alice hides her maternal alleles. The hiding generates ***M***, the variant alleles of the metacohort. In step 2, the metacohort variant alleles are sent through the untrusted channel. In step 3, the untrusted computes disease risk for each metasample and generates the disease risk for all of the L metasamples. The results are sent back to Alice in step 4. After receiving the risk for all the metasamples, Alice discards the risk estimates for all the metasamples except for her own. ***b)*** Variant selection procedure in projection-based (1) and allele sampling-based hiding (2). In projection-based hiding, ***I*** is compared to generation panel ***G*** and closest individual is identified. Next, the variants whose alleles match between ***I*** and the alleles of closest individual are selected. In sampling-based hiding, all the variants are used. ***c)*** Projection-based hiding. The variants are selected first by comparison to ***G***. After variant selection, ***G*** is sampled with replacement to generate the metasamples. Finally, Alice is added to the metasamples to generate the metacohort. ***d)*** Allele sampling-based hiding. The variants alleles are sampled using the allele frequencies estimated from ***G*** (Step 1). After sampling, the metacohort variant alleles are generated for L-1 metasamples. Finally, Alice’s alleles are inserted among the other metasamples. Alice’s alleles are inserted in the metacohort as the Meta-2’ sample in a random position that is uniformly distributed among the metacohort.

Alice first hides her alleles using HaploHide and generates the hidden variant alleles dataset (denoted by ***M***), which we refer to as a metacohort, with her hidden alleles. The metacohort contains the alleles of *m* variants for *L* individuals. We refer to the samples in the metacohort as metasamples. Alice’s alleles are hidden among the metasamples, which essentially represent the “crowd” in which Alice’s data is hiding. Alice sends the matrix that contains the metacohort alleles to the other party (Fig. 1). The other party performs computation for each metasample in the metacohort and sends the disease risk estimates in an list back to Alice. Alice discards the results for all the individuals but herself and uses her disease risk estimate. The hiding performed by HaploHide enhances Alice’s privacy because her variant alleles are not directly distinguishable from the variant alleles of metasamples. In this scenario, Alice sends the alleles of one *haplotype*, i.e. maternal or paternal copy of her variants. The other haplotype must also be hidden in another independently generated metacohort and sent to the untrusted party. After Alice receives the results for both haplotypes, she combines the results of disease risk estimates from two haplotypes (See Combination of Haplotype-Specific Results).

### Hiding Framework and the HaploHide Algorithm

HaploHide uses two approaches to hide variants depending on the correlation of the alleles among the variants. The first approach, we refer to as “projection-based hiding”, is used when there is high inter-variant correlation. The variants whose alleles are highly correlated are generally inherited together as haplotypes in strong linkage disequilibrium (LD) blocks[39]. In general, it is known that the genomic distance between variants correlate with the strength of LD blocks. When the variants are in LD or in haplotype blocks, the hiding must be performed by respecting the LD blocks such that the untrusted party cannot utilize the correlation between the variants to identify Alice in the metacohort. In order to correctly simulate haplotypes, HaploHide utilizes a panel of individuals as a metacohort generation panel (denoted by ***G***). In projection-based hiding, the Alice’s alleles are “projected” onto ***G***. This operation finds the individual in ***G*** whose alleles are closest in distance to Alice’s alleles (Fig. 1b). Since Alice’s alleles may not match exactly to one of the individuals in ***G***, this hiding approach may decrease the number of variants that are hidden. However, we observed that when variants are in strong LD blocks, a close match can be found and almost all the variants can be hidden (“Projection-based Hiding”). The projection-based hiding is effective on variants that are located over short genomic distances such as variants on individual genes.

After selection of variants, HaploHide generates the metacohort by generating the metasample variant alleles. Figure 1c illustrates the generation of metacohort with projection-based hiding. In projection-based hiding, Alice’s shared alleles match exactly to one of the individuals in ***G***. The other metasamples’ variant alleles are generated by sampling random individuals from ***G*** (Fig 1c). Thus, the metacohort is exactly a subsample of individuals from ***G***. The motivation behind this is to ensure that the metacohort alleles reflect the exact haplotype structure of the variants in ***G***. We demonstrate the use of this approach for hiding of the variants on genes.

The second approach is used when the variants are not in strong LD blocks, i.e. far from each other in genomic coordinates, and the inter-variant allele correlation is low. We refer to this approach as “allele sampling-based hiding” or simply as “sampling-based hiding”. In this case, HaploHide uses ***G*** to estimate the allele frequency distribution of the variants. These distributions are then sampled for generating the variant alleles for each metasample in the metacohort. Figure 1d shows the generation of the metacohort using sampling-based hiding. For each metasample, variant alleles are generated independent of other variants. For each variant, HaploHide samples an allele from the distribution of alleles for the corresponding variant. The allele distribution for each variant is estimated from ***G***. Since the correlation among variant alleles is low, the alleles are independently sampled for each variant. We demonstrate the use of this approach for hiding GWAS variants that are spread over a long genomic region. In comparison, projection-based hiding aims at conserving the haplotype structure of the variants while sampling-based hiding aims to use the fact that the inter-variant allele correlation is low and conserve only the allelic frequencies.

### Importance of Generation Panel on Privacy Provided by Hiding-based Framework

A vital component of effective hiding is how diverse the generation panel is and how well the generation panel matches to the ancestry of the individuals that are being hidden. When the generation panel ***G*** matches the ancestral population of the hidden individuals then the generated metacohorts are very similar, in statistical properties, to the randomly generated sets of samples from ***G***. Thus, without any auxiliary information, a metacohort of size *N* generated by HaploHide looks very similar to a random sample of size *N* generated from ***G*** and the adversary cannot learn anything from the metacohort more than what can be learned from a random sample generated from ***G***.

It must also be noted that this condition is generally not trivially satisfied because every genetic ancestry must be fairly represented in ***G***, which may not be accomplished in practice. In addition, ***G*** may not be representative of all of the individuals whose haplotypes are being hidden. Therefore, the generated metacohorts may be biased more towards certain populations. As the genomic panels are diversified, we foresee that ***G*** will be more representative of the genetic ancestries of majority of the individuals.

#### Leakage Statistics and Identification Attacks on Metacohorts

We analyze the re-identification attacks under practical scenarios in order to evaluate the extent to which the hidden individual is identifiable under changing parameters of hiding such as changing ***G*** and changing variant-variant correlations. The identification attacks on the metacohort involves the untrusted party (or adversary) to try and identify the hidden individual (for example, Alice) in the metacohort. We first consider the ways that the untrusted party can attack the hidden allele datasets to recover Alice’s alleles. For this, the adversary must use a scoring function to sort the metasamples and reveal the metasample with highest (or lowest) scoring metasample as the identified hidden individual (Fig. 2a). To simulate the attack procedure, we compute a number of statistics, which we call *leakage statistics* (See Methods), to characterize how effective hiding is. The leakage statistics represent the features that adversary uses to identify the hidden individual within the metacohort. For each metasample, the leakage statistics are computed and the distribution of each statistic is generated. We then evaluate whether the hidden individual is at the extremes of this distribution.

**Figure 2.**
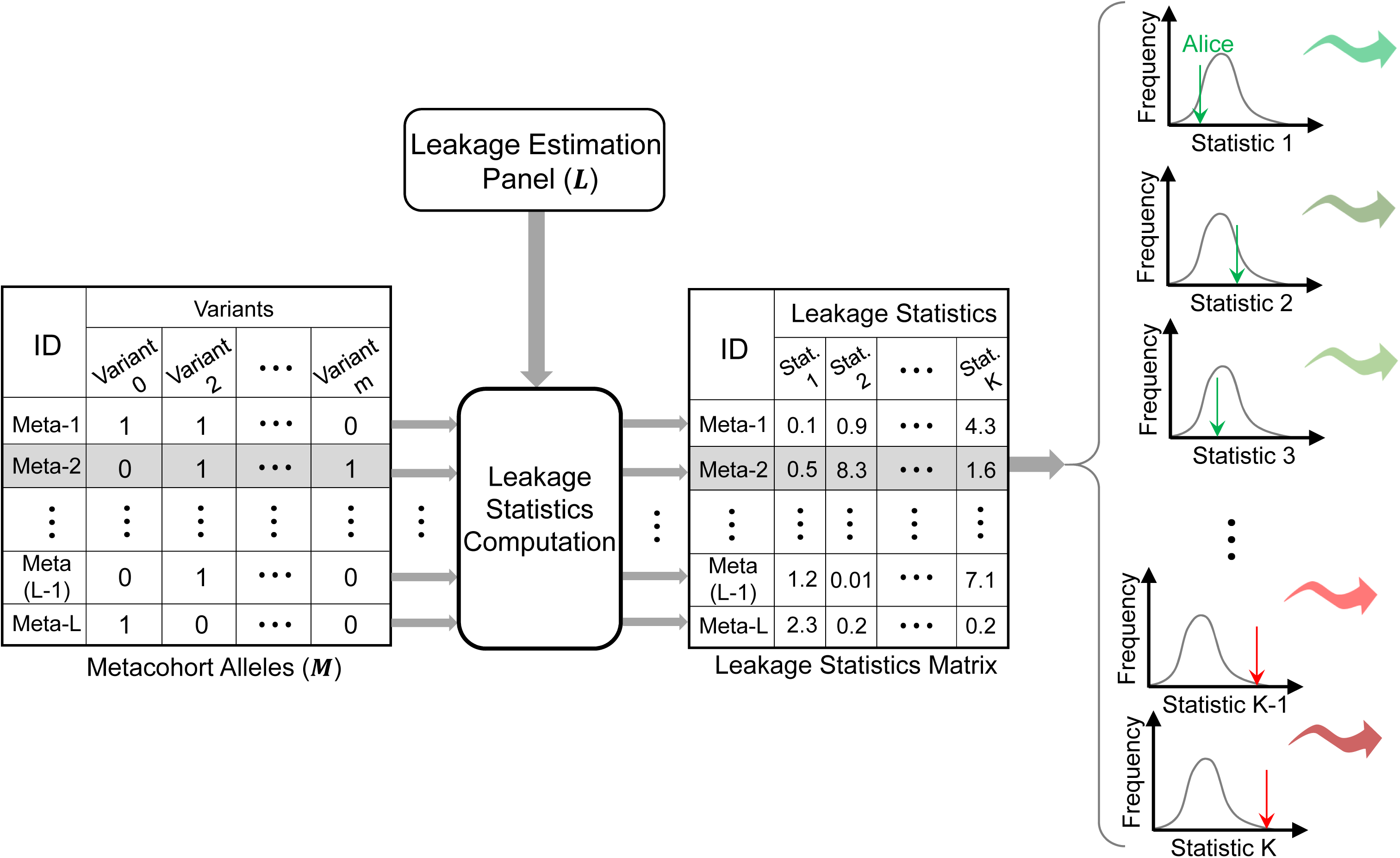
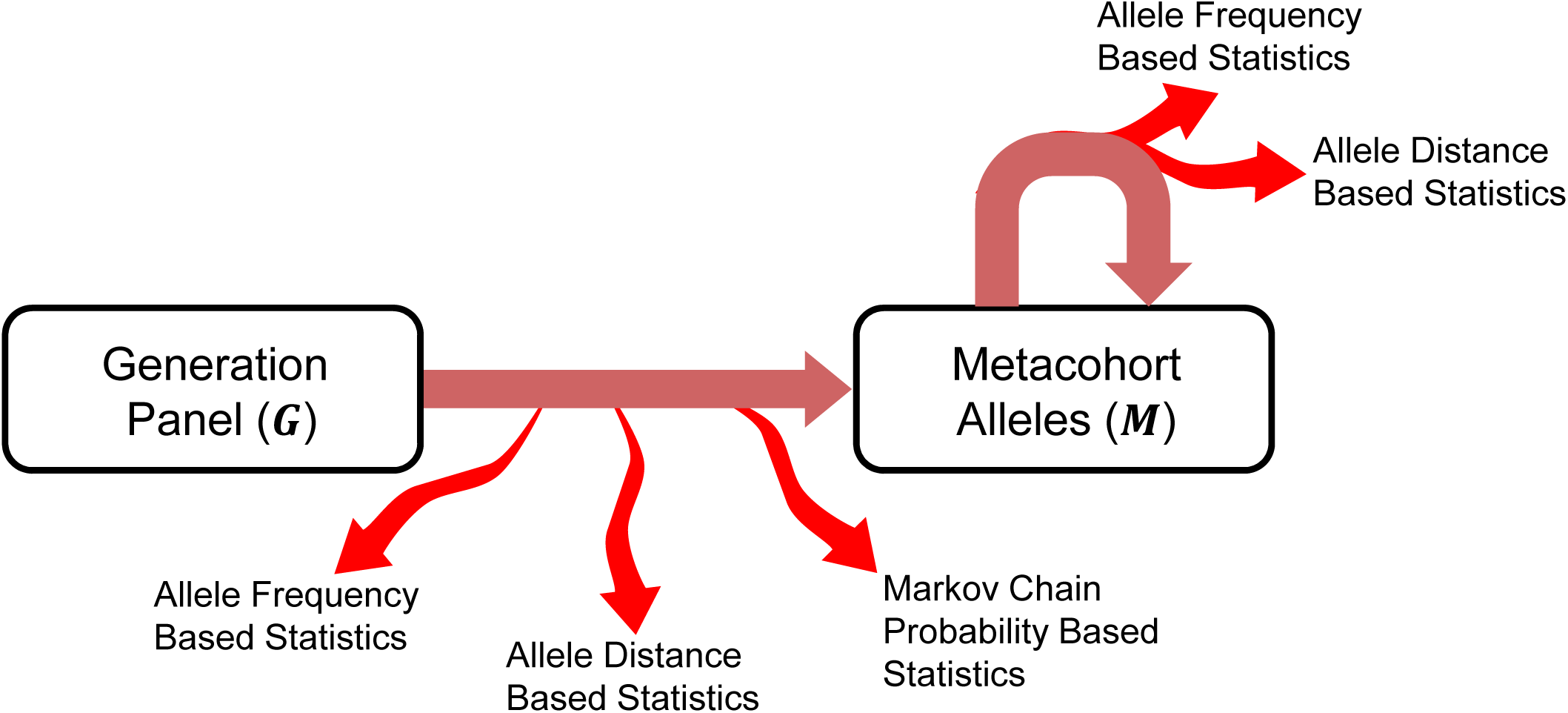
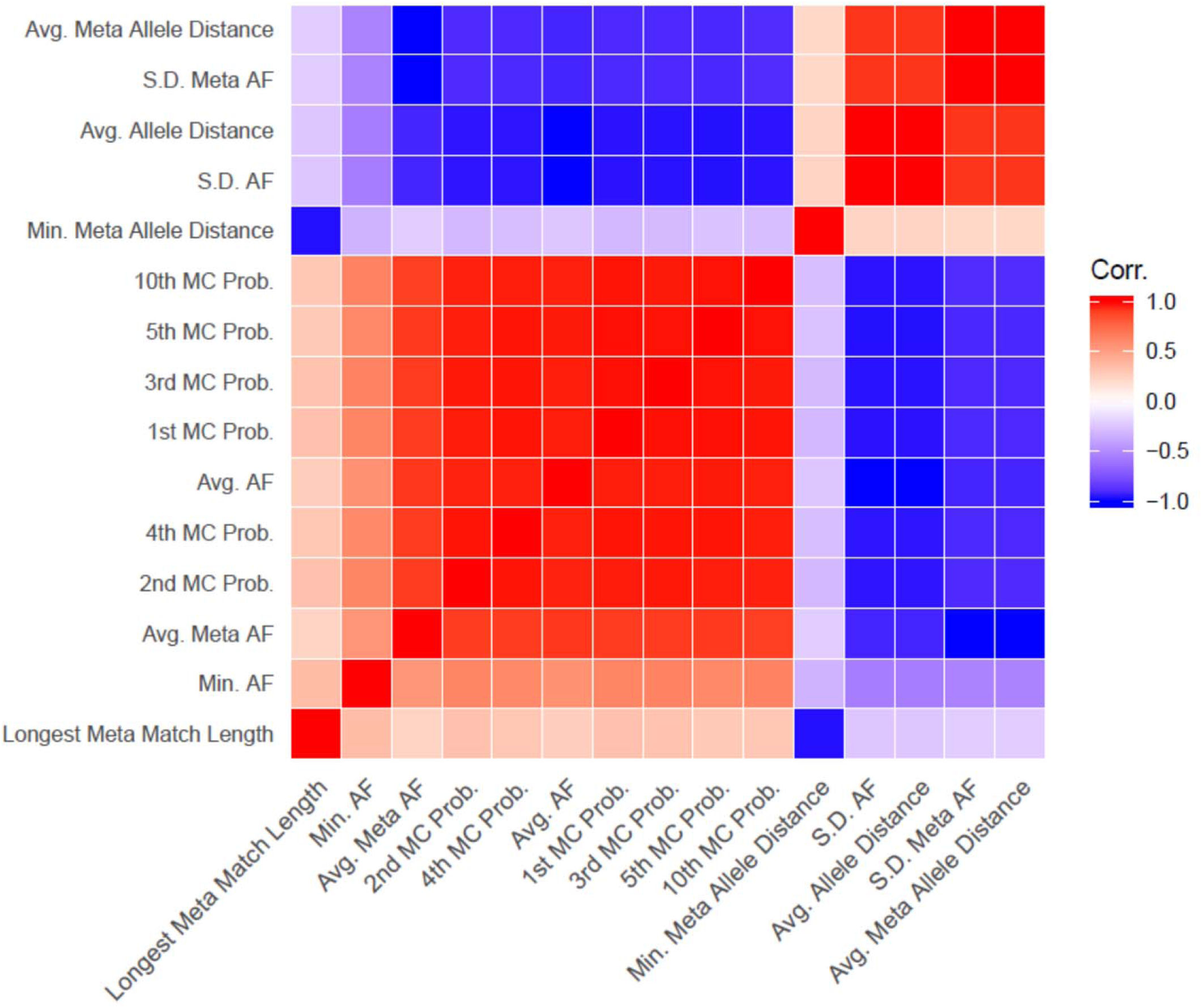
Illustration of leakage statistics. ***a)*** Leakage statistic matrix computation. The metacohort variant alleles is used to compute K statistics for each metasample. Alice’s rank among metasamples for each statistic is computed to assess identification of Alice among metasamples. For this, the distribution of each of the K statistics among metasamples is computed and Alice’s rank is identified, as illustrated on the right. The statistics for which Alice is at the extremes represent statistics that an adversary can use for re-identification. ***b)*** The illustration of the types of leakages from hiding framework. Three types of leakage statistics are illustrated. ***c)*** Correlation matrix among different leakage statistics as illustrated by a heatmap. High and low values are shown with red and blue colored cells. The statistics are referenced by abbreviations. See Methods for description of the abbreviations.

The leakage statistics have been compiled by reference to the previous studies that have shown what type of statistics lead to accurate individual re-identification[40]. It is worth noting that these statistics are certainly not the complete list of leakage statistics that an adversary could use. In fact, it is not possible to enumerate every possible statistic that an adversary could use. Rather, these statistics represent the set of features that were the most widely demonstrated for performing re-identification attacks.

While computing the leakage statistics, we assume that the adversary makes use of an external panel of variant genotypes. Adversary uses this panel to learn the properties of the variants, such as allele frequencies and the LD blocks. We refer to this panel as the leakage estimation panel and denote it by ***L*** (Fig. 2a). ***L*** and ***G*** are considered as analogs in hiding and in re-identification attack because ***G*** is used as a knowledge-base for hiding individuals and ***L*** is used by the adversary as a knowledge-base to identify individuals. We assume that Alice will chose ***G*** to be equal to the population panel that corresponds to the ancestral population of herself, which we denote by ***A***. Similarly, we assume that adversary aims to maximize identification accuracy by choosing ***L*** to be equal ***G*** (or ***A***). We evaluate how the leakage statistics get affected when ***L*** and ***G*** mismatch between each other and between ***A*** (Section “Mismatchesbetween Generation and Leakage Estimation Panels”).

Figure 2b illustrates the different types of leakage of information that can be used to identify hidden individuals in a metacohort. We divide the statistics into 3 main categories: Allele distance-based statistics, allele frequency-based statistics, and Markov chain probability-based statistics.

##### Allele Distance Based Statistics

Computation of distances between the variant alleles has been used in linking attacks for identifying individuals between independent datasets. The basic motivation is to identify the individual in one dataset whose alleles match best to the individuals in the other dataset. A good match is used to link these individuals in two datasets. In the hiding scenario, the allele distance-based leakage statistics are used to directly compare the leakage panel individuals to the metasamples to reveal whether any metasample is substantially similar (or unsimilar) to the leakage panel individuals. For instance, the metasample with the largest distance to the leakage panel individuals (farthest neighbor of leakage panel individuals) may identify the hidden individual.

##### Allele Frequency Based Statistics

It has been discussed previously that the allele frequency of the variants can be used for identification purposes. For example, the existence of many rare alleles can be very identifying in linking attacks. To account for the allele frequency-based attacks, we compute, for each metasample, the minimum and maximum allele frequency among the variants. The allele frequency-based statistics measure the allelic composition of the variant alleles. For example, the metasamples with high number of low-frequency variants may be re-identify the hidden individual.

##### Markov Chain Probability Based Statistics

Several previous studies showed that the correlation among variants is a very strong predictor and can be used to identify individuals in linking attacks[41], [42]. For this, Markov chains are used to model the dependence between the alleles of consecutive variants. As the dependence of alleles manifest over the haplotype blocks, it is necessary to account for the correlation among the alleles of variant that are consecutive over long stretches. This is shown in previous studies where high order Markov chain probabilities can increase the power of identification. We evaluate multiple orders of Markov chain probability estimates for evaluating re-identification.

If the statistic computed from Alice’s alleles is an outlier within the distribution of statistic values among the metasamples, Alice can be easily pointed out within the metacohort. In this case the hiding is not successful. However, while Alice may be extreme for a statistic, this does not necessarily mean that the statistic is an effective statistic for identifying everyone because the statistic may not be extreme for other individuals. Therefore, it is necessary to evaluate whether each leakage statistic can *systematically identify individuals* at a large scale. To test this, we perform hiding of individuals and evaluate leakage statistics for each individual. We analyze the distribution of the leakage statistics and assess whether any statistic or combination of statistics can systematically identify individuals in practical scenarios. We also evaluate whether the joint usage of leakage statistics can be used to systematically identify hidden individuals.

We assume that the attacker does not have access to auxiliary information. The main reason for this is that if the adversary already knows the variants genotype, it would be non-sensical for the adversary to identify the hidden individual within the metacohort to learn her variant genotypes. If the attacker gains access to auxiliary genetic information, the attacker needs to have access to at least *log_2_*(*N*) bits of identifying genetic information to correctly identify an individual within a metacohort of *N* individuals. The auxiliary identifying information can be stolen/hacked or acquired by other means such as prediction from gene expression. In principle, the risk of auxiliary information can be remedied by increasing the metacohort sample size.

We first evaluated the relationship between the leakage statistics. For this computation, we generated metacohort for hiding the CEU samples in the 1000 Genomes Project using the European samples (EUR super-population) as the metacohort generating population. We next computed the leakage statistics for each metacohort and computed the pairwise correlation matrix of the statistics. Figure 2c shows the correlation between the different statistics that are assigned to the individuals in the metacohort. The statistics exhibit substantial correlation between each other. This indicates that although we are evaluating a repertoire of leakage statistics that represent a broad spectrum of leakage sources, none of them provide orthogonal information independent of the other statistics. This suggests that most of the identifying information from these statistics is highly redundant and may not bring extra re-identification power to the adversary.

### Projection-based Hiding: Variants on Cancer Predisposition Genes

We demonstrate the use of projection-based hiding of variants for hiding the cancer-gene variants. This scenario is summarized in Fig. 3a. In this scenario, Alice would like to send her variant alleles on a gene that is associated with cancer risk. The variants that Alice sends include her germline variants and a number of variants that are only specific to Alice. These variants can be de-novo or extremely rare variants. These variants are important for evaluating the cancer risk because they generally have an unknown impact, i.e., similar to variants of unknown significance. The risk estimation aims at analyzing all the variants on the cancer predisposition genes and assessing the cancer risk of Alice.

**Figure 3.**
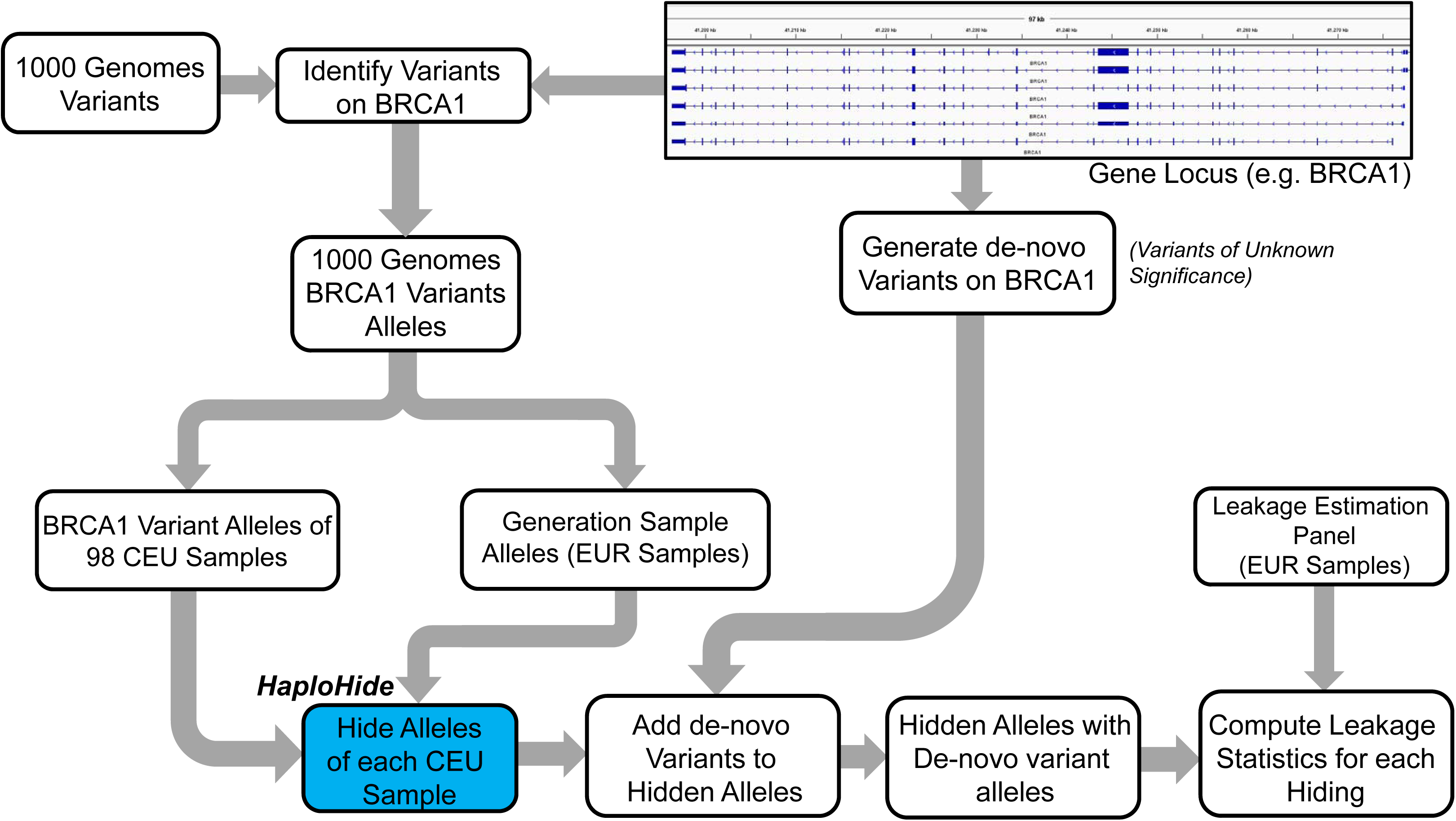
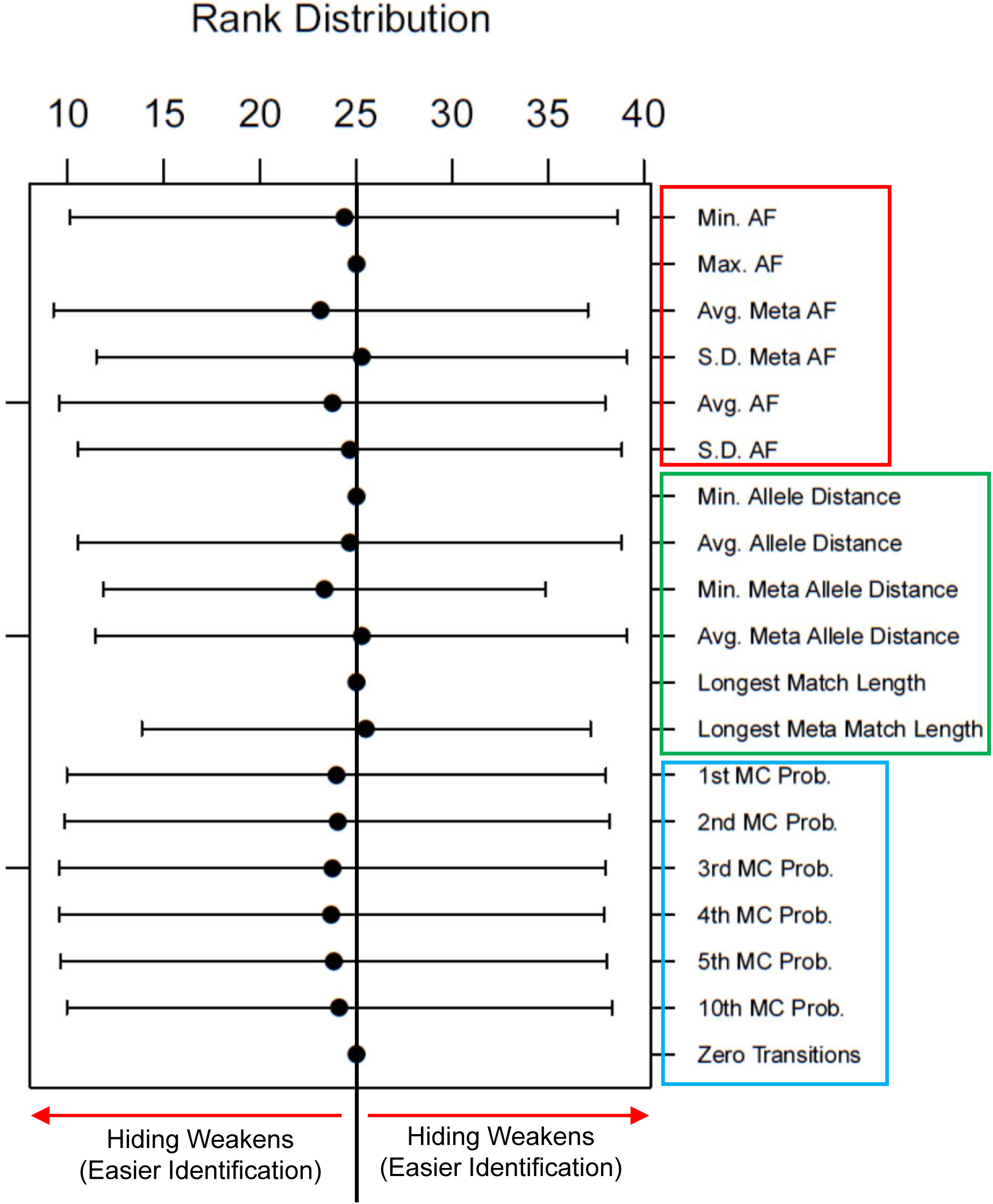
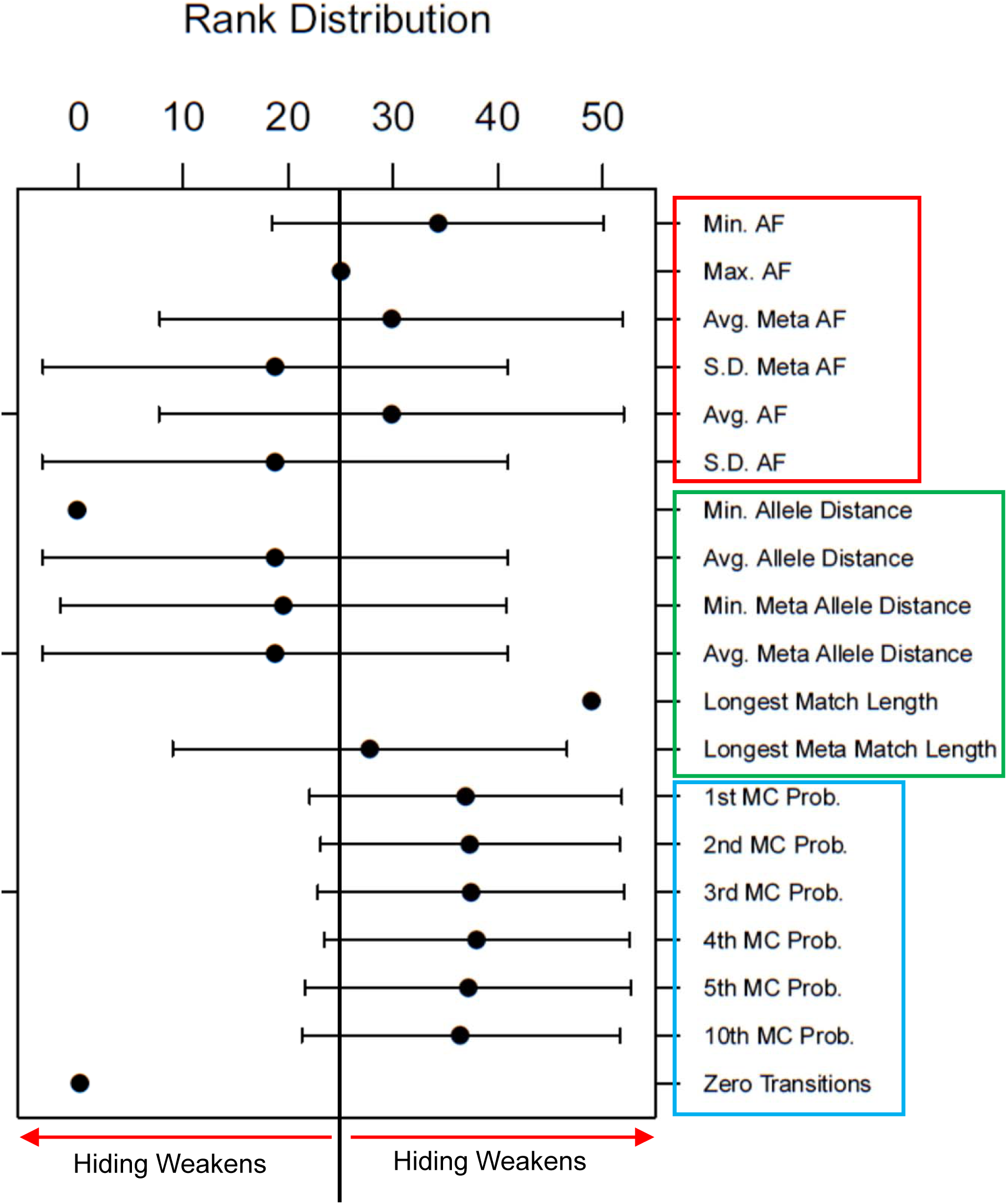
***a)*** Illustration of the projection-based hiding for sharing variant alleles on cancer predisposition genes. The variant alleles on the gene of interest (BRCA1 in the figure) is extracted for the 98 CEU samples sequenced in The 1000 Genomes project. 98 CEU samples are used as the hidden individuals. The individuals of European ancestry (EUR) in The 1000 Genomes Project are used as the generation panel. For each of the 98 CEU samples, a metacohort is generated. After generation of the metacohort, the de-novo variants are introduced into the metacohort. After the de-novo variants are introduced, the leakage statistics are computed for each hiding using the EUR samples as the leakage estimation panel. ***b)*** The distribution of leakage statistic ranks (y-axis) of the CEU samples for each statistic (x-axis). Each bar represents the rank distribution. The mean rank is shown with the dot. The error bars represent one standard deviation. Different statistics are grouped together with rectangles where allele frequency-based statistics are grouped with red box, allele distance-based statistics are grouped with green box, and Markov chain-based statistics are grouped with a blue box. The middle rank (25) is marked at the middle with a line. ***c)*** The leakage statistic rank distributions using the allele sampling-based hiding of the variant alleles on cancer predisposition genes.

Alice hides her alleles using projection-based hiding to generate the metacohort variant alleles (***M***). In the hiding, de-novo variants are introduced to each metasample (Fig. 3a, Methods Section). This way we simulate the existence of de-novo variants in the genomes of metasamples. After the generation of the metacohort, Alice sends the metacohort alleles to the company that will perform risk assessment for all the metasamples in the metacohort. The results are then sent back to Alice where she discards all the results except for her own.

In order to perform a general evaluation of the leakage statistics, we used the 98 individuals of Central European descent (CEU) in the 1000 Genomes Project, i.e. ***A*** is the set of individuals in Central European descent. We focused on BRCA1, BRCA2, and PTEN genes that are associated with risk of breast cancer. For each individual, the variants on these genes with higher than 1% minor allele frequency are extracted. Next, we generated the metacohort using the European samples (EUR super population) as the metacohort generation panel (***G***), using projection-based hiding. As a comparison we also generated the metacohort using random allele sampling-based hiding. Using this procedure, we generated 196 metacohorts (98 using projection-based, 98 using sampling-based hiding) for each gene. Each metacohort contains 51 samples such that there are 50 metasamples and 1 hidden individual. We finally computed the leakage statistics for each metacohort using the European samples as the leakage estimation panel (***L***). The current set of leakage statistics do not evaluate the leakages from the de-novo variants and therefore we assume that the de-novo variants do not leak any information to re-identify the hidden individual. We deem that this is a reasonable assumption because the distribution of de-novo variants within small distances, i.e., regions smaller than 1 megabases, do not show a specific non-uniform distribution and are almost uniformly distributed [43]–[46]. On another note, the number of de-novo variants per sample is very small and it is unlikely that the de-novo variants can be used to re-identify the hidden individual when the attacker does not have access to auxiliary information.

To visualize the distribution of leakage statistics for each individual, we computed the rank of the hidden individual among the metasamples with respect to each leakage statistic. This enables us to evaluate whether the hidden individual is an extreme with respect to the leakage statistics. Figure 3b illustrates the distribution of ranks of the hidden individual with respect to the leakage statistics when we use projection-based hiding. An important observation is that all of the statistics show an approximately symmetric distribution around the mid-rank (rank of 25). To simplify visualization of large number of statistics, we divided the statistics into different classes that we introduced previously. This result highlights that hidden individuals are spread effectively among the metasamples with respect to the leakage statistics when projection-based hiding is used.

Projection-based hiding performs a variant selection step where the variant alleles of the individual are compared to every sample’s alleles in the generation panel and the best matching sample’s matching variants are used in generating the metacohort (Fig. 1b, Methods Section). This operation may decrease the number of variants that is hidden in the metacohort. In our tests, we observed that more than 99.9% of the gene variants (60,646 out of 60,685 total variants in all samples) can be shared when projection-based hiding is applied on variants that are in strong LD blocks.

We next applied sampling-based hiding on the cancer gene variants and computed the distribution of leakage statistic ranks for hidden individual. This is illustrated in Fig. 3c. We observed that there are clear deviations from the mid-rank for several statistics. For example, all of the Markov Chain probability-based statistics deviate to higher ranks. This result shows the fact that real individual is always at higher ranks for these statistics when sampling-based hiding is used to generate the metacohorts. Thus, an adversary can utilize Markov Chain probability-based statistics to identify the hidden individual in the metacohorts when sampling-based generation is used. Moreover, other types of statistics may be used to identify the hidden individual. For example, the hidden individual is systematically located at higher or lower ranks for several allele frequency-based statistics.

Consequently, our results provide initial evidence that the projection-based hiding can be practically feasible using a population panel as small as the 1000 Genomes Project for hiding variant alleles on genes where the variants are in strong LD blocks, i.e., variants that are distributed within short distances. In addition, projection-based hiding retains virtually all of the variants. In contrast, sampling-based hiding cannot effectively hide the variants in strong LD blocks as it does not respect the correlative structure between alleles of close-by variants.

### Mismatches between Generation and Leakage Estimation Panels

In the previous scenario, we used generation panel (***G***) and leakage estimation panel (***L***) to be close to the ancestral population panel (not exactly equal) of the hidden individuals (***A***) in terms of haplotype structure and frequencies. This is necessary to ensure probabilistically safe hiding (“Comparison of The Probability Distribution of Metacohorts” in Methods Section and Supplementary Information). An important consideration in hiding is how the mismatches between generation panel, leakage estimation panel, and the ancestry of hidden individual affect the hideability of individuals. In particular, it is useful to measure how the leakages change when generation panel does not exactly match to the ancestral population of the hidden individual.

We tested how a mismatch between ***G***, ***L***, and ***A*** would affect hiding and leakage statistics. For this, we focused on mismatches between the populations of generation and leakage panels as illustrated in Fig. 4a. For comparison, we included 2 other super-populations from the 1000 Genomes Project, namely American (AMR) and African (AFR) in addition to the European populations (EUR). We used the individuals in these populations as ***G*** and ***L*** while generating the metacohort that hide the CEU samples. The ranks distributions of leakage statistics when (***G*** ≈ ***A***) are shown in top row in Fig. 4b. It can be seen that the real individual’s ranks are generally distributed symmetrically around the mid-rank, which indicates that the hiding is effective when the hidden individual’s population matches the generation panel population.

**Figure 4.**
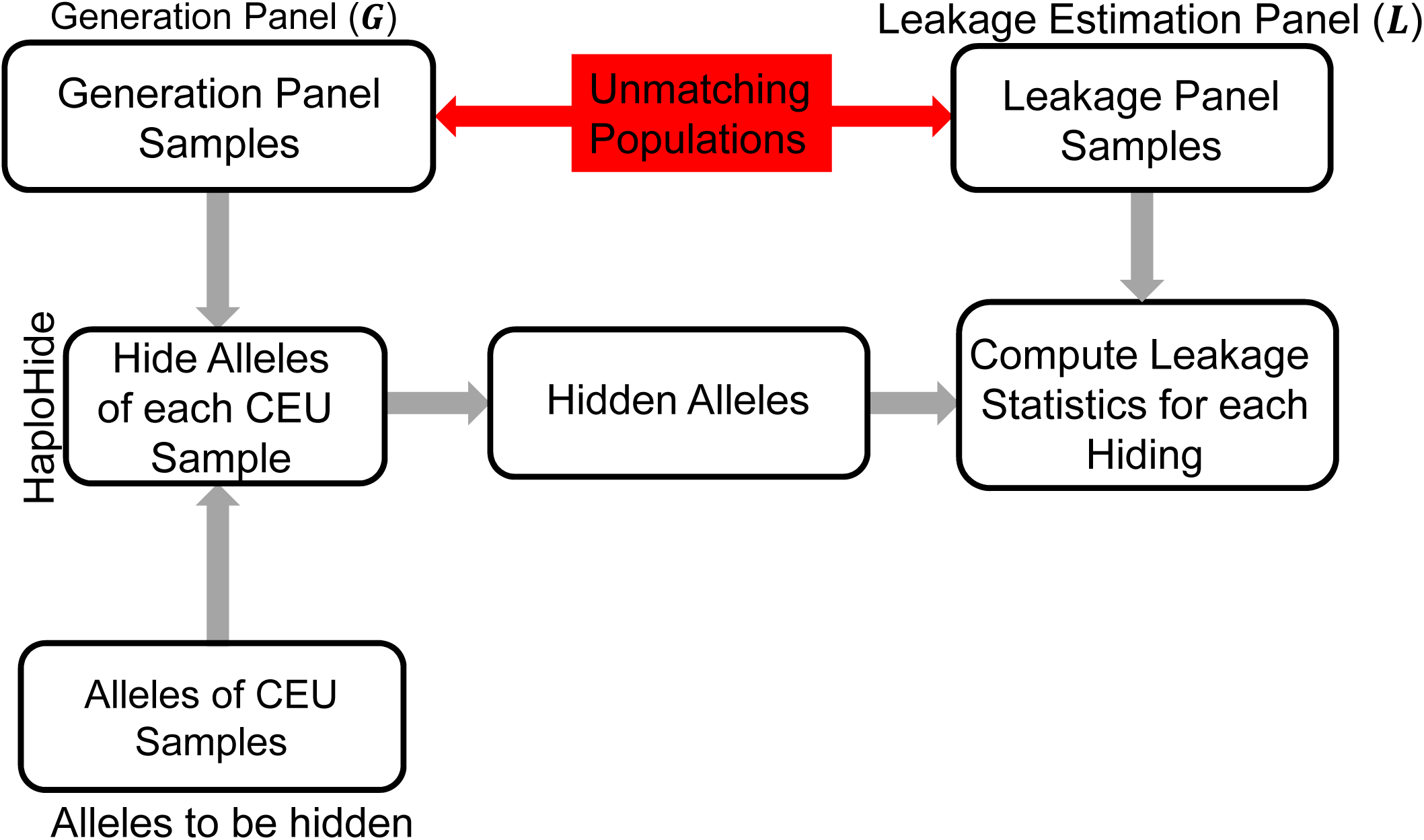
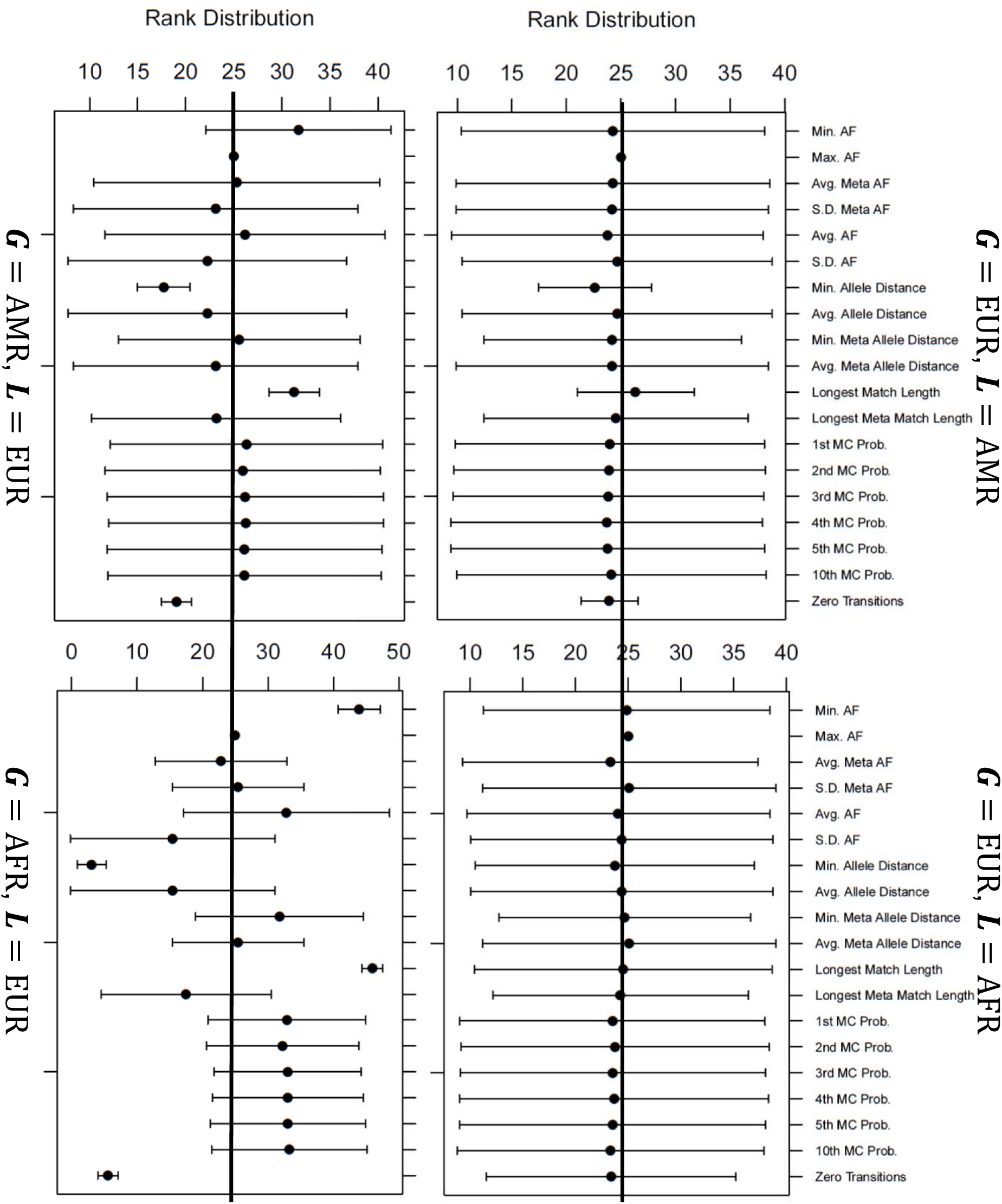
***a)*** The population mismatches between the generation panel, ***G***, and the leakage estimation panel, ***L***. The 98 CEU samples are hidden using HaploHide with alternative ***G*** and ***L***. ***b)*** The leakage statistic rank distribution computed with ***G*** and ***L*** alternating between AMR, AFR, and EUR. Top row shows the distributions for ***G*** = *EUR* and ***L*** = {*EUR, AMR*}. Bottom row shows the distributions for ***L*** = *EUR* and ***G*** = {*EUR, AMR*}.

Interestingly, when ***G*** = EUR and ***L*** = AMR, we did observe two of the allele-distance based statistics shifting slightly (Fig. 4b). This can be attributed to the fact that the CEU samples show similarities to the American populations because American populations contain admixture of European and Native American populations[47]. This highlights the importance of matching the genetic background of hidden individuals to the generation panel in hiding-based framework. Nevertheless, the deviation of these statistics is not substantial and increasing the metacohort size can help in blending of the hidden individual in the crowd.

Next, to evaluate the effect of generation panel population unmatching the hidden individual’s population, we used AMR and AFR samples as generation panel and used EUR samples as the leakage estimation panel (***G*** ≠ ***A***, ***L*** ≈ ***A***). The distributions of leakage statistic ranks are shown in bottom row in Fig. 4b. It can be seen that many statistic ranks show a substantial deviation from the mid-ranks, i.e., ineffective hiding. This is especially observable when AFR samples are used for generation and EUR samples are used for leakage estimation.

These results show that it is vital to ensure that the generation panel population matches the hidden individual’s population (***G*** ≈ ***A***). This follows the condition that we derived for probabilistically safe hiding (Methods Section) and it is expected since the generation panel is used to generate the crowd among which we are hiding the hidden individual. If the population of hidden individual is different compared to the crowd, it is fairly easy to identify this individual within the crowd because the hidden individual’s genetic information is easily detectable among a non-matching crowd. Our results show the very important role of genetic background of Alice and building a “good crowd” for effective hiding.

### Joint Usage of the Leakage Statistics

Up until now, we treated each leakage statistic independent of each other. In reality, an adversary can use the leakage statistics jointly for identification. This is illustrated in Fig. 5a. The adversary computes the leakage statistics for each metasample and generates a “leakage statistics matrix”. The adversary then uses the matrix to identify the hidden individual. To visualize whether the joint usage of statistics will reveal any pattern that can be exploited in an identification attack, we computed the leakage statistics for all metasamples and performed dimensionality reduction on the leakage statistics matrix using the t-distributed stochastic neighbor embedding (tSNE), which is an extremely effective method for non-linear dimensionality reduction[48]. We used tSNE algorithm to decrease the dimensionality of the leakage feature matrix (Methods Section) and evaluate whether the small mismatches in ***G*** and ***A*** can be detected.

**Figure 5.**
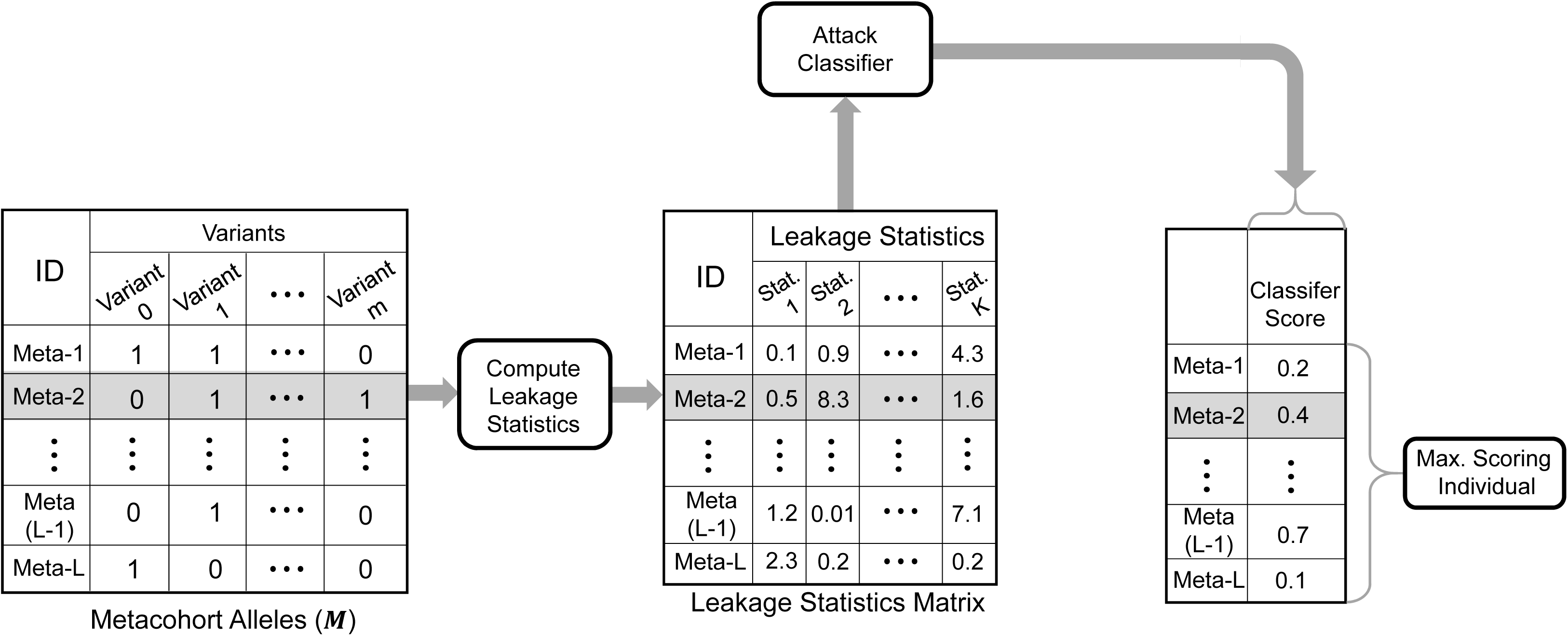
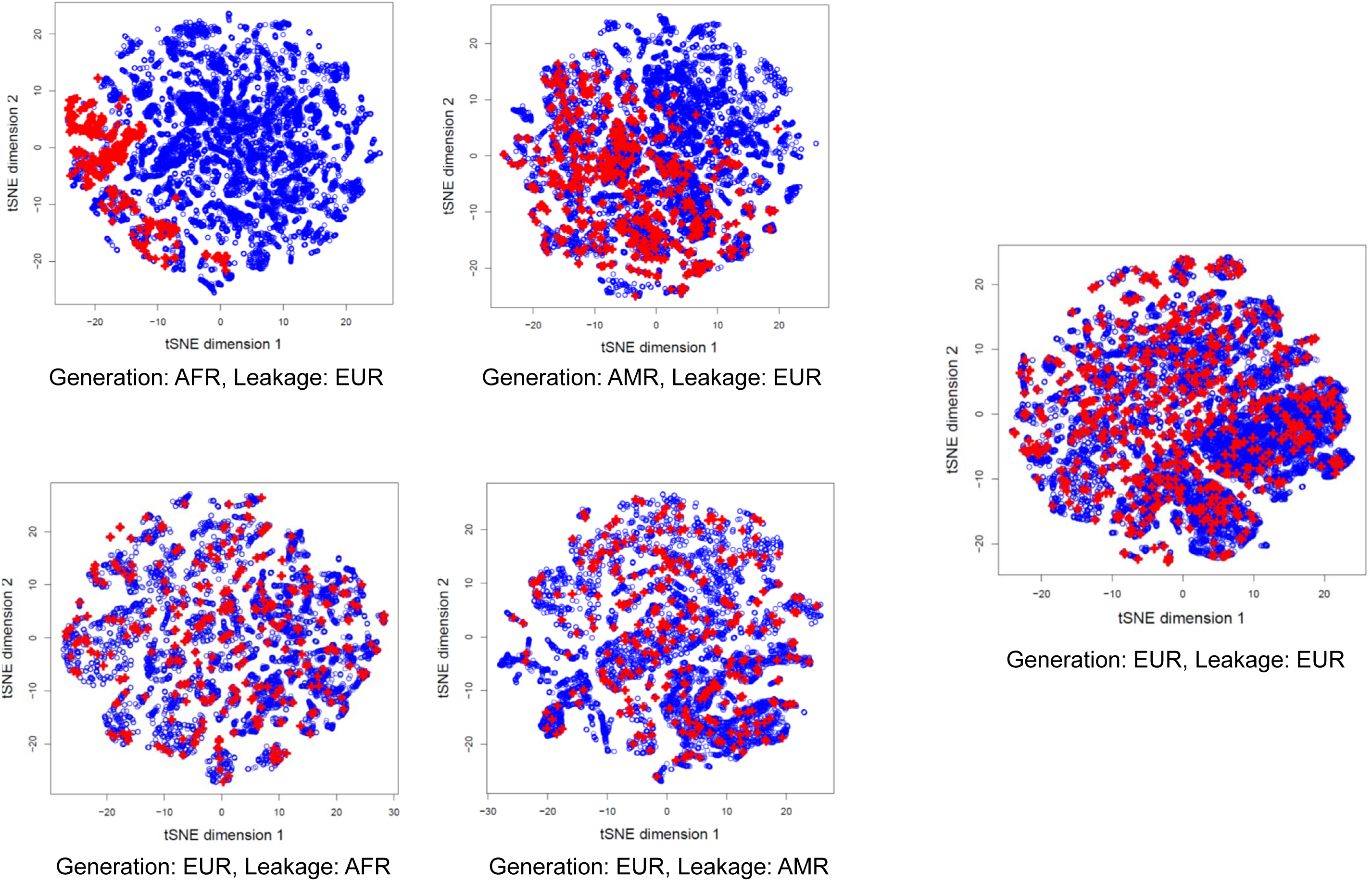
***a)*** Illustration of the adversarial attack. After the leakage statistics matrix is computed, the matrix is used in building an attack classifier that combines all the leakage statistics to make use of the features jointly to identify the max-scoring individual as the hidden individual. ***b)*** Scatter plots show the tSNE-based dimensionality reduction of the leakage statistic matrices. Each dot in the scatter plots represent the a metasample. The real hidden individuals are indicated by red dots on the plots. Each scatter plot corresponds to the leakage distributions computed with different generation and leakage estimation panels, as specified by labels below each scatter plot.

Figure 5b shows the tSNE plots for different options in selection of generation and leakage estimation panels. It is clearly seen that the hidden individuals (red points) are clustered in specific locations in the dimensionality reduced data when the generation panel’s population does not match the hidden individual’s population. In contrast, when the hidden individual’s ancestral population matches generation panel’s population (***G*** ≈ ***A***), the real individual’s data is distributed without a clear pattern within the low dimensionality data. These results suggest that even the joint usage of the features is not systematically useful for identifying the hidden individuals.

### Effect of Linkage Disequilibrium

We next studied the effect of linkage disequilibrium on projection-based hiding. It is generally known that the linkage disequilibrium is a function of distance between variants. To test this effect, we first randomly picked 10 regions in the genome. Next, we identified 50 variants in each region such that the variant-variant distance is at least Δ*d* base pairs and we changed Δ*d* = {1000, 10000, 50000} (Fig. 6a). This way, for each Δ*d*, we generated a set of variants that show a different level of LD among variants (Fig. 6b). We finally performed projection-based hiding of the CEU samples for the selected variants and computed the leakage statistics. We used EUR samples as both the generation and leakage estimation panels. Figure 6c shows the rank distributions for each set of samples that are identified by the corresponding Δ*d*. It can be seen that hiding is effective when min (Δ*d*) is larger than 1000 base pairs but it is not effective for min(Δ*d*) > 10,000 bps. This result shows that while projection-based hiding is effective for hiding the variants in strong LD blocks (i.e., gene regions), it starts failing for hiding variant alleles in low LD blocks, i.e., variants whose alleles are independent from each other.

**Figure 6.**
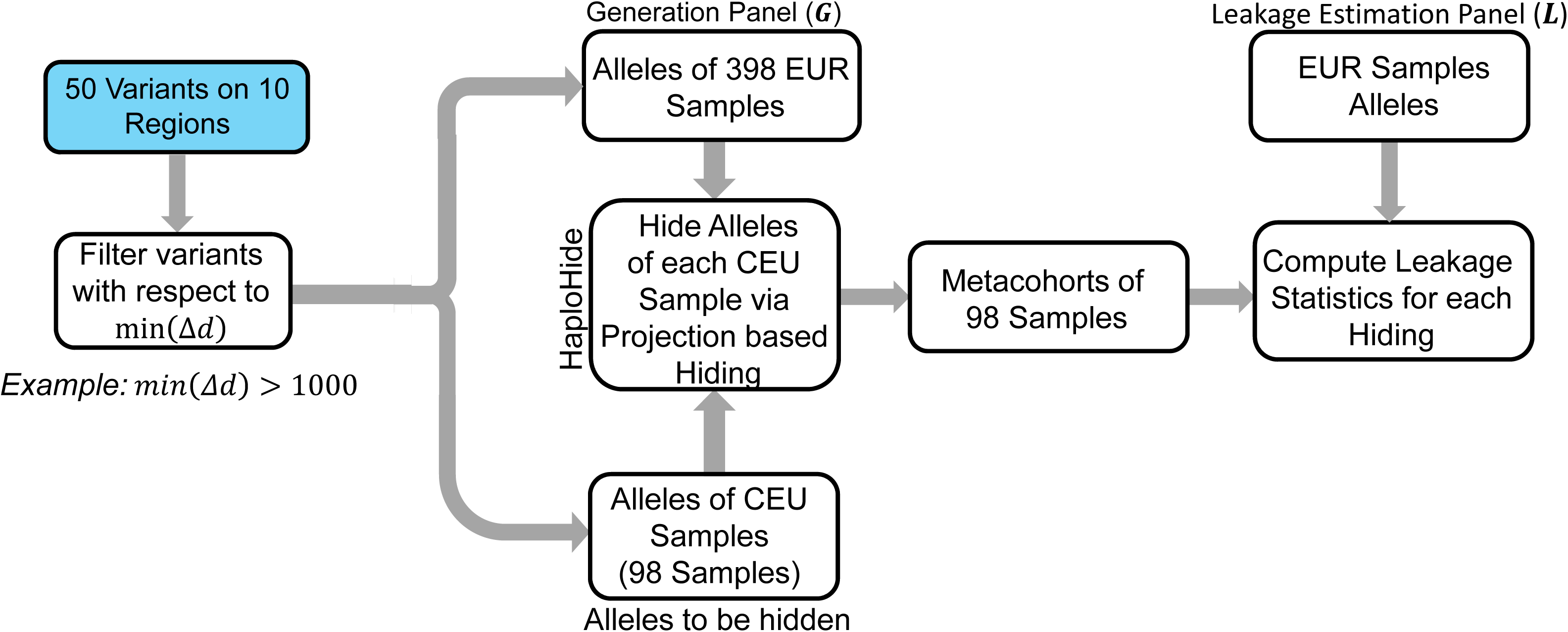
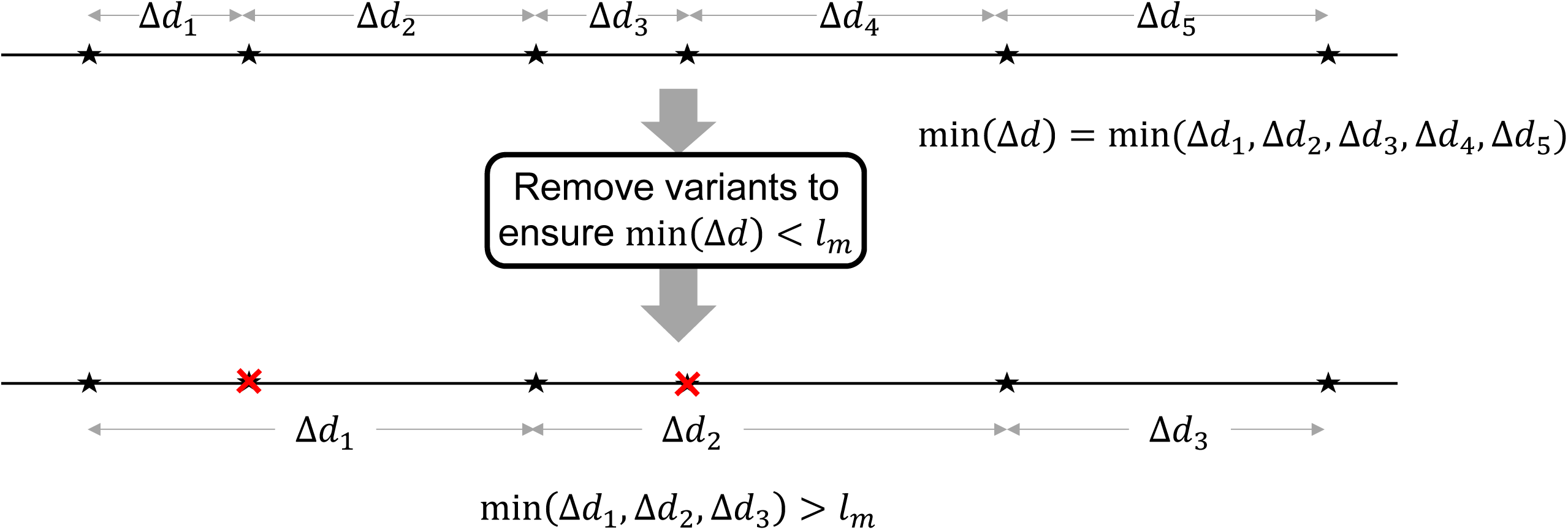
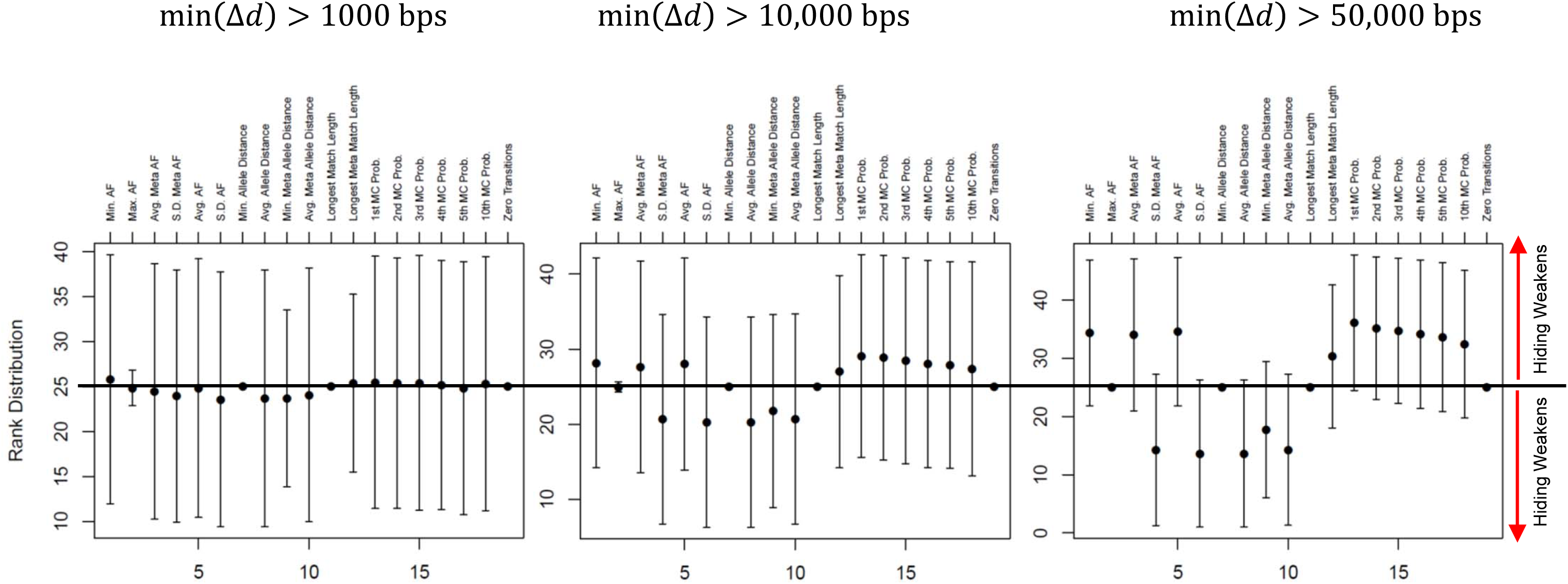
***a)*** The effect of changing variant-variant distances on the hiding framework. CEU samples are hidden using variants after they are filtered with respect to min(Δ*d*) distance. ***b)*** min(Δ*d*) distance constraint among the list of variants. ***c)*** Distribution of leakage statistic rank of the hidden individual with changing min(Δ*d*) constraint, as indicated above the plots.

### Sampling-based Hiding: GWAS Variants

We next focus on the usage of sampling-based hiding approach for hiding variant alleles in low LD regions. One of the potential applications of hiding weak or no LD variants is sharing of variants identified by genome-wide association studies (GWAS). The GWAS variants are found to be significantly associated with existence of a certain trait or phenotype and they are scattered all over the genome. Therefore, most of these variants are generally not under influence of linkage disequilibrium. This is reasonable because the variants that are in strong LD blocks do not effectively provide new information with respect to how much they contribute to the genotype-phenotype associations. These independent variants alleles can generate the constitutive component of the polygenic risk scores.

The GWAS variant hiding scenario is summarized in Fig. 7a. For demonstrating the hiding of GWAS variants, we first extracted the variants that are associated with cardiovascular diseases from GWAS catalog on chromosome 1, which yields 44 variants. Then we filtered the variants such that min(Δ*d*) > 1,000,000 bps and identified 33 variants that satisfy this criterion. This filtering ensures that there is no linkage disequilibrium among the selected GWAS variants. We next used allele sampling-based hiding to generate the metacohort (using 50 metasamples and 1 hidden sample) to hide the GWAS variant alleles of the CEU individuals. We used EUR samples as generation panel. For each individual’s metacohort, we computed the leakage statistics using EUR samples as leakage estimation panel. Figure 7b shows the rank distribution of leakage statistics for the hidden individuals. The ranks of hidden individuals are symmetrically distributed around 25. This indicates that the sampling-based hiding can be used to effectively hide the variants that are under weak or no linkage equilibrium blocks. In comparison with the projection-based hiding, sampling-based hiding complements the hiding framework for hiding of the variants such that there is weak or no LD among the variants.

**Figure 7.**
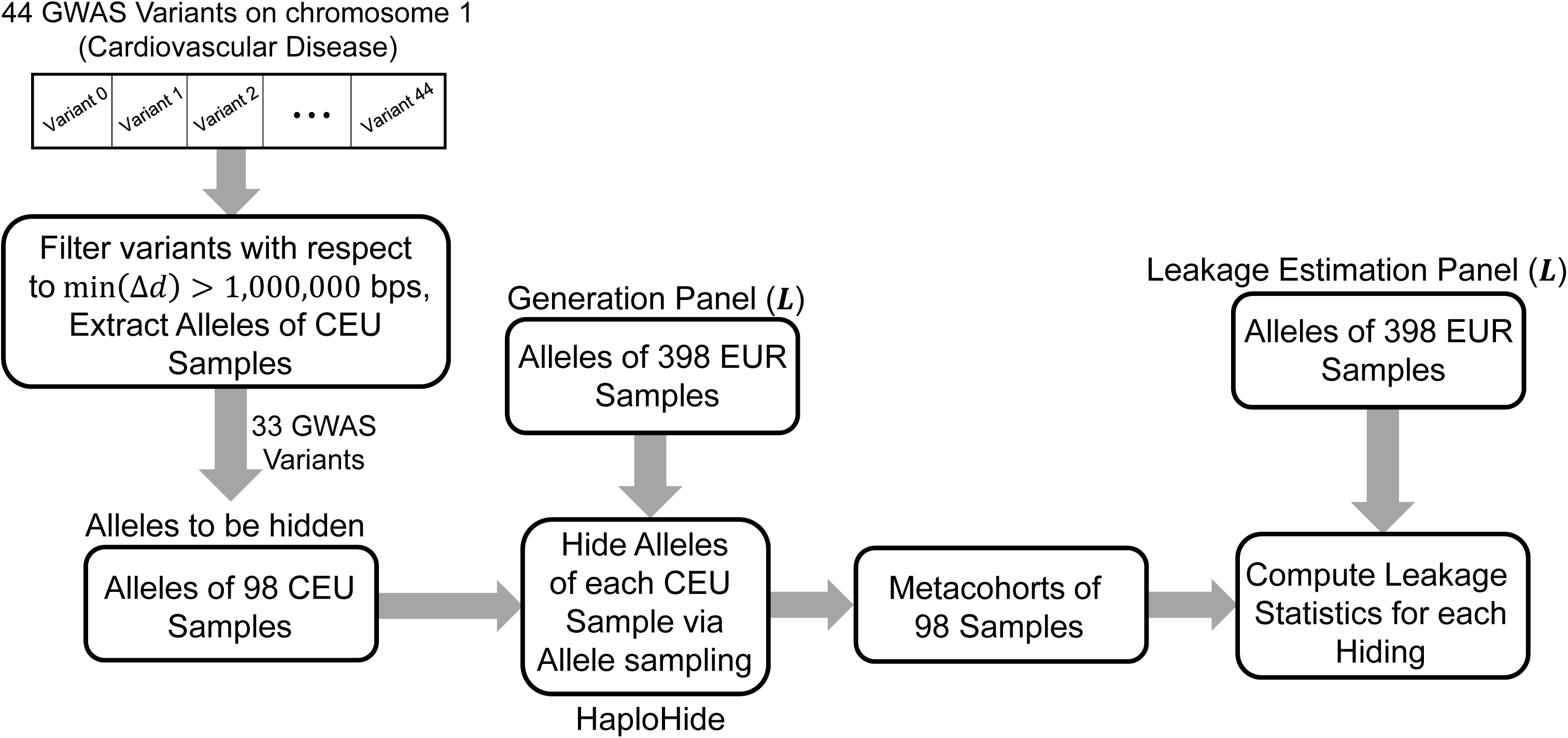
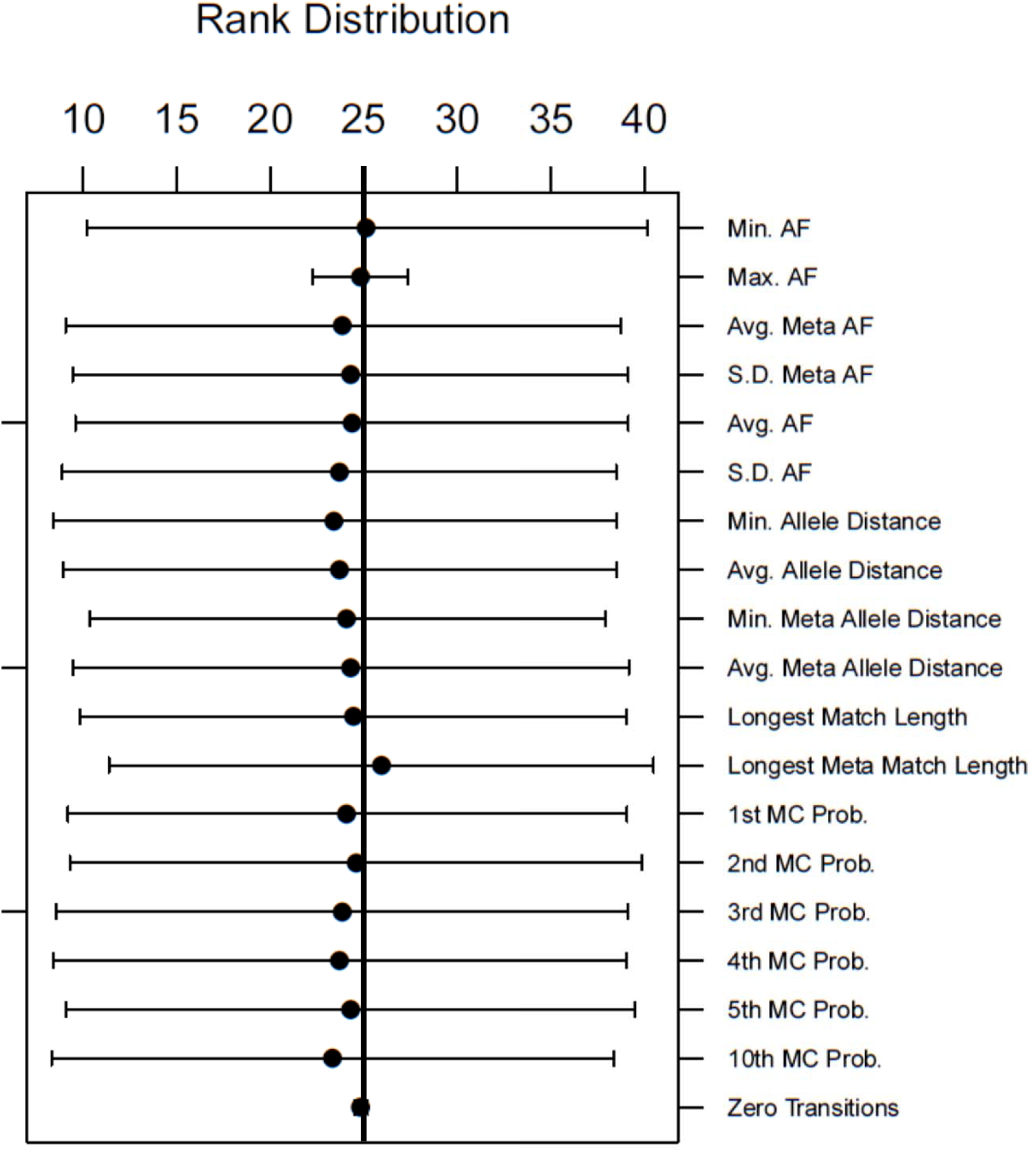
GWAS variant hiding statistics. ***a)*** The GWAS variant allele hiding using allele sampling-based hiding. The GWAS variants are filtered with respect to min(Δ*d*) > 1,000,000 bps, which yields 33 variants. Next, alleles of 98 CEU samples are extracted. Next, the variants are hidden using allele sampling. In hiding, EUR samples are used as the metacohort generation sample. Finally, the leakage statistics are computed using EUR samples as the leakage estimation panel. ***b)*** The distribution of leakage statistic rank for the hidden individuals.

### Combination of Haplotype-Specific Results

In the currently presented scenarios, Alice must separate her genotypes into two haplotypes (maternal and paternal), hide the variant alleles in the two haplotypes independently (i.e., run HaploHide twice) then send the 2 metacohorts to the untrusted party (Fig. 8) separately. After the company (untrusted party) performs risk estimation on each haplotype, the results will be returned to Alice independently. Alice then discards, for each metacohort, the results for the metasamples except her own result then combine the result from paternal and maternal haplotypes into the final disease risk estimate. The need for accurately combining the results from maternal and paternal haplotypes may limit the applicability of the hiding-based framework. However, many disease risk estimation methods (such as polygenic risk scores[14]) can be decomposed in terms of the risk estimated for each haplotype. For example, most of the polygenic risk score estimation methods are based on linear combination of the risk alleles weighted by their odd-ratios. In this regard, the risk estimates from maternal and paternal haplotypes can be simply combined by a multiplication or summation. Since the combination operation can be performed by Alice, it does not incur any increase in the privacy risk. It should be noted, however, that there are newer approaches for risk estimate that are based on very large number of risk alleles using complex computational approaches[49]. The current hiding framework does not accommodate these approaches.

**Figure 8.**
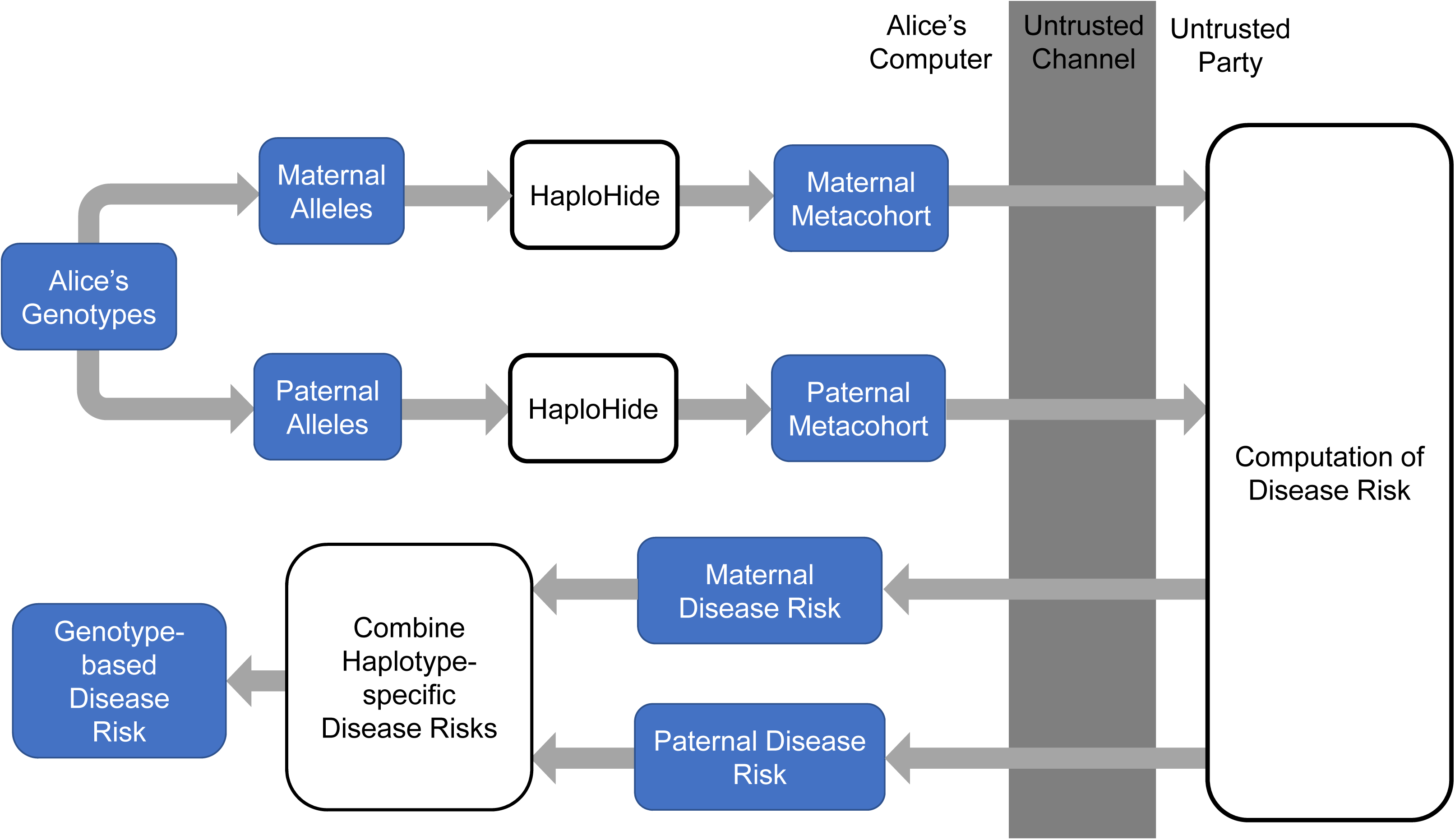
Block diagram illustrates the combination of the disease risk using the allele specific results. The genotypes are first separated into paternal and maternal haplotypes. Next, each haplotype is hidden independently using HaploHide. The metacohorts for the maternal and paternal haplotypes are sent to the untrusted party for estimation of disease risk. The disease risk estimation for maternal and paternal haplotype are sent back to Alice independently. These risks are combined by Alice to generate the genotype-based disease risk.

## Discussion

In this paper, we present HaploHide, a novel genomic data hiding framework for privacy-enhanced personal genomic data sharing. HaploHide aims at hiding the variant alleles in a metacohort such that the hidden individual’s alleles are hidden in a crowd of metasamples. Conceptually, HaploHide has a simple motivation that is based on protecting privacy by hiding-in-the-crowd or a camouflaging strategy: If Alice utilizes a generation panel that is representative of her ancestry and genetic background, she can hide her genetic information effectively such that an adversary cannot systematically identify her variant alleles among the crowd of metasamples. Based on this simple conceptual idea, HaploHide represents the first proof-of-concept approach for assessing the feasibility of hiding based frameworks for genomic data. The genomic hiding can be implemented using more complicated approaches. For example, hiding methods based on synthetic haplotype generation algorithms[50] can enable more effective hiding by modeling of the haplotype structures. It is, however, not clear whether the haplotypes generated by these tools can be directly used for hiding genomic data with guarantees on probabilistically safe hiding. This represents a future outlook for hiding-based frameworks.

We foresee that a very important ingredient in hiding based framework is to evaluate the leakages that may enable the adversary to identify the hidden individual. To study leakages, we proposed a large set of statistics that can be used to identify the hidden individual. These statistics comprehensively describe the allelic composition of the metasamples to evaluate whether the hidden individual is located at an extreme of any of the statistics. Thus, the leakage quantification must be an integral part of the hiding process so as to evaluate whether the hiding is effectively performed and to ensure that the generation panel is effective in hiding the individual’s alleles.

The basic premise of data hiding strategy depends on the diversity of the metacohort generation panel. This panel is used as a knowledge-base of haplotypic structure for realistically generating the metacohort in which the individual’s variant alleles are hidden. We demonstrated that the population mismatch between the individual to be hidden and the generation panel may cause ineffective hiding. Thus, existence of extensive generation panels is imperative for performing effective hiding. The population scale sequencing projects[3], [5], [7], [51], [52] that generate data for detailed delineation and mapping of genomic diversity in different populations will greatly benefit developing effective generation panels for safely hiding of individuals from minority populations. The publicly available portions of these datasets will enable us to build extensive metacohort generation panels including thousands of samples from diverse set of populations. These cohorts will also enable characterization of a large spectrum of variants, such as very-rare variants.

In the hiding, we demonstrated the usage of two use cases where two different approaches are used for hiding. The first approach, we termed projection-based hiding generates a metacohort by projecting the hidden individual’s variant alleles on the generation panel samples. We showed that this approach is effective for hiding the alleles of variants that are in strong LD blocks. In the second approach, HaploHide utilizes an allele sampling procedure. This approach is effective when the variants show very weak or no correlation with each other. These two scenarios represent two extreme cases haplotypic structure: The very strong LD blocks where projection-based hiding is effective and very weak LD blocks where sampling-based hiding is effective. A potential future question is hiding variants in a mixture of strong and weak LD blocks. In these cases, new hiding methods must be developed to appropriately model the complex haplotype structure and generate realistic metacohorts where we can hide variant alleles. Larger generation panels will enable modeling these complex haplotypes better. This point reinforces the importance of using a diverse generation panels in hiding. We have highlighted that the importance of matching between the generation panel and the genetic background of hidden individuals in qualitative terms and not in strictly quantitative terms. A more formal approach can be used to analyze and quantify the upper and lower bounds on identifiability when the generation panel does not match to the genetic backgrounds of the hidden individuals.

Compared to other formalisms such as differential privacy[28] and cryptographic approaches, e.g., homomorphic encryption[31] and multiparty computation[32], the advantage of hiding-based framework is that it does not require a fundamental change in the way data is processed. It only requires a lot more data processing. For example, in encryption-based data sharing, both parties must perform homomorphic computations and this imposes fundamental changes in the implementations of the underlying algorithms. Moreover, it imposes heavy increases in the required computational resources. In comparison to these approaches, data hiding framework does not require an increase in the computational complexity. In principle, the untrusted party does not even have to change the way that they perform their computations on the genomic data. They only need to perform computation on every metasample in the metacohort. Thus, the hiding framework can be immediately deployed with minimal overhead on the existing infrastructure. Most notably, the data hiding idea is related conceptually the k-anonymization, its variants, and extensions [53]–[56]. In summary, k-anonymization aims at sanitizing a database (such as a table) such that no combination of columns in the table can identify less than k individuals in the table. In this sense, k-anonymization aims at sanitizing and protecting every individual within the database. This is an inherently different problem than hiding that we employ in the current study, where we aim at protecting the privacy of an individual’s privacy, i.e. sample size of 1.

## Dataset Availability

The phased genotypes and the population information are obtained from The 1000 Genomes Project[2] at https://ftp-trace.ncbi.nih.gov/1000genomes/ftp/release/20130502/.

GWAS variants are extracted from the GWAS Catalog[57].

## Online Methods

We describe the hiding mechanisms, leakage statistics, and analytical underpinnings of the hiding framework.

### HaploHide Algorithm

HaploHide algorithm takes as input the variant alleles of Alice, denoted by ***I*** and the metacohort generation panel, ***G***. ***I*** contains the alleles for *m* variants that Alice will share with the untrusted entity. Each allele is a binary value {0, 1} where 1 indicates the existence of alternate allele at the respective variant and 0 indicates the reference allele. The generation panel, ***G***, contains the alleles of the *m* variants for *N* individuals, i.e. binary matrix with *N* rows and *m* columns. The generation panel is a database that is representative of the haplotype structure and the allele frequency distribution of the variants. The output from HaploHide is the metacohort variant alleles, which we denote by ***M***. This is a matrix that contains *m* variant alleles for *L* metasamples. The variant alleles of Alice are included in one of the rows of ***M***. ***M*** is sent to the untrusted entity.

After the untrusted party receives ***M***, they will analyze each metasample in ***M*** and return the result for each metasample to Alice. In the re-identification, untrusted party analyzes ***M*** to identify which alleles belong to Alice. While performing re-identification, we assume that the adversarial party uses another external panel, which we refer to as re-identification panel, denoted by ***L***. Leakage panel is independent of the generation panel but we assumed that the adversary knows the generation panel exactly and can use it as re-identification panel, i.e. ***L*** = ***G***.

#### Projection-based Hiding Mechanism

There are 3 steps in the projection-based hiding. First is the projection of Alice’s alleles on the generation panel to identify the closest generation panel individual. Second step is variant selection. Third step is random sampling from the generation panel to build the metasample set.

##### Projection of Alice’s Variant Alleles

HaploHide first identifies the individual in ***G*** which has the smallest Hamming distance in terms of allele difference to Alice, which we refer to as the projection step:

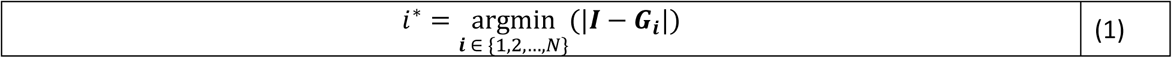

where *i*^∗^ denotes the index of the individual in the generation panel whose alleles are closest to Alice’s alleles, ***I***.

##### Variant Selection

After the projection, the variants are filtered such only the variants matching variants are selected:

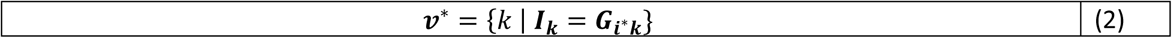

where ***v***^∗^ denotes the set of *t* indices within [1, *m*] for variants whose alleles match between ***I*** and ***G***_***i***_. Then the matching variant alleles of this individual are shared. Finally, the alleles for the selected variants are used. The variant selection ensures that the non-matching variant alleles of the projected individual are not shared.

##### Random Sampling of Generation Panel

Finally, ***G*** is randomly sampled with replacement *L* times to generate the metasamples. At each sampling, the alleles of variants at the indices ***v***^∗^ are extracted for each sampled individual.

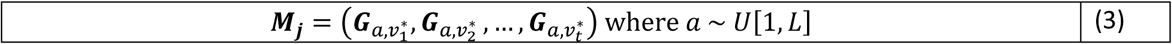

where ***M***_j_ denotes the alleles of *j*^*th*^ metasample. Finally, the alleles of variants at indices ***v***^∗^ are extracted from ***I*** and randomly placed within the metacohort by selecting a uniformly distributed position among the metasamples. *U*[1, *L*] denotes the uniform distribution of integers in [1, *L*].

##### Introduction of de-novo variants

We introduced de-novo variants to the cancer predisposition genes by random uniform sampling of genomic positions on the genes.

#### Allele Sampling-based Hiding Mechanism

In allele sampling-based hiding, HaploHide first computes the allele frequencies of alleles in the *m* variants in ***I***;

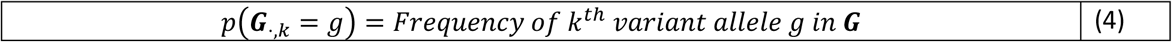

Next, for each metasample, the allele frequencies are sampled to generate the variant alleles:

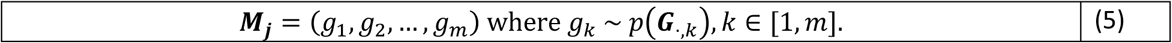

Thus, the allele frequencies are independently sampled for each variant (and for each metasample). Finally, Alice’s variant alleles are randomly placed within the metacohort.

### Comparison of The Metacohorts with Randomly Generated Samples from ***G***

We qualitatively compare the samples generated by HaploHide and by random sampling. In this comparison, we will lay out the necessary conditions for ensuring that the probability of generating a sample by hiding is the same as generating the same sample by random sampling of ***G***. This is important as it implies that an adversary cannot learn anything from the metacohort more than what he/she can learn from a random sample of individuals from ***G***. This way, the hiding does not bias certain metacohorts compared to random sampling. In other words, if the samples (i.e. metacohorts) generated by HaploHide look exactly like random samples from ***G***, the hiding can be deemed effective.

We assume that we are given the set of variants (***v***), the generation panel ***G***, and the ancestral population panel ***A*** of the individuals to be hidden. The variants specify the loci for which haplotypes are shared. It should also be noted that we constrain the set of hidden individuals (and haplotypes) to be in the panel ***A***. In this setup, we discuss the conditions for which the following probabilities are equal:

a. Probability of generating a set of haplotypes (denoted by ***Q***) by random sampling from ***G***
b. Probability of generating ***Q*** as the metacohort while hiding a randomly selected individual from ***A***.

We formulate this as:

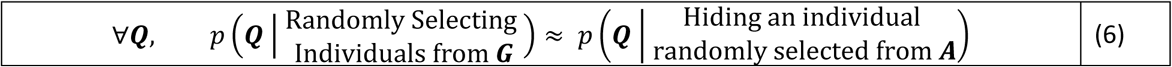

where ***Q*** denotes a sample of haplotypes of *L* individuals from ***G***. In (6), 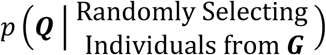 denotes the probability of randomly selecting the *L* individuals (or haplotypes) from ***G*** to generate ***Q***. In this case, we assume that each individual is uniformly randomly selected from ***G***.

In (6), 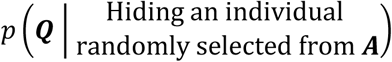 denotes the probability of generating ***Q*** as the metacohort of *L* individuals as we hide an individual that is randomly selected from ***A***. We use ***G*** as the metacohort generation panel while hiding. For simplicity of discussion, we focus on the projection-based hiding. In this setup, hiding is a three-step process:

1. We first randomly select an individual from ***A***
2. Next, we project the selected individual onto ***G***, then select the closest individual. We assume that the projection does not reject any variants, i.e., there is a perfect match for any individual in ***A***, in terms of haplotype comparisons.
3. Finally, we select (*L* − 1) samples randomly from ***G***.

From step 3, *L* − 1 individuals are generated by random sampling from ***G***. Thus, the only difference between random selection of individuals from ***G*** (left hand side in (6)), and the projection-based hiding (right hand side in (6)) is the generation of the hidden individual. More specifically, the difference is the random selection of the hidden individual from ***A*** followed by the projection on ***G*** (Supplementary Figure 2) in the hiding. To be concise, we refer to sampling from ***A*** followed by the projection on ***G*** as “sample from ***A*** then project to ***G***” and we refer to sampling from ***G*** as “sample from ***G*** directly”. Below we discuss the general condition that “sample from ***A*** then project to ***G***” and “sample from ***G*** directly” are identical.

Sampling from ***A*** generates a sample with respect to the distribution of the haplotypes in ***A***. The projection maps the individual (or haplotype) sampled from ***A*** to an individual in ***G*** with respect to distance between haplotypes, i.e., the sample from ***A*** is mapped to an individual in ***G*** with respect to maximal haplotype match. If each and every individual in ***A*** can be matched uniquely to a distinct individual in ***G*** such that that every individual in ***G*** is a projection of exactly one individual in ***A*** (projection is one-to-one and onto), it is straightforward to see that “sample from ***A*** then project to ***G***” is equivalent to “sample from ***G*** directly”. However, projection may not be one-to-one or it may not be onto. For example, some individuals in ***G*** may be selected more (or less) frequently than other individuals by the projection just because their haplotypes are rarer (or more frequent) in ***G*** compared to ***A***. Then the projection may create a bias by selecting some individuals more (or less). For example, some individuals in ***A*** may not have any good haplotype matches in ***G*** and we may not be able to project them on ***G***. In this case “sample from ***A*** then project to ***G***” deviates from “sample from ***G*** directly”. Generalizing on this observation, if the frequencies of unique haplotypes of individuals in ***A*** are similar to the frequencies of the unique haplotypes of individuals in ***G***, then “sample from ***A*** then project to ***G***” will be very similar to “sample from ***G*** directly” (Supplementary Figure 2a). In essence, this condition implies that any haplotype in ***G*** (and ***A***) must be represented in ***A*** (and ***G***) with similar frequencies. If similar haplotypes in ***A*** and in ***G*** have similar frequencies, the projection will not distort the haplotype frequencies and therefore “sample from ***A*** then project to ***G***” will have the same distribution as “sample from ***G*** directly”. This condition further implies that if the genetic background of individuals, in terms of haplotype frequencies, in ***G*** and ***A*** are similar, then “sampling from ***A*** then project to ***G***” is essentially same as “sampling from ***G*** directly”.

### Computation of Leakage Statistics

We present the details of how leakage statistics are computed.

#### Allele Distance Based Statistics

Allele distance-based statistics are computed by comparison of the variant alleles between the metacohort and the re-identification panel, ***L***. These statistics are used by the adversary to identify the hidden individual (i.e., Alice) among the metacohort samples. We list these statistics below:

##### Average and Minimum Allele Distance Statistic

This statistic measures the average distance between the metasample variant alleles and the leakage panel sample variant alleles. We present several metrics that are based on comparison of alleles between samples in leakage panel and the metacohort. The allele distance is defined as:

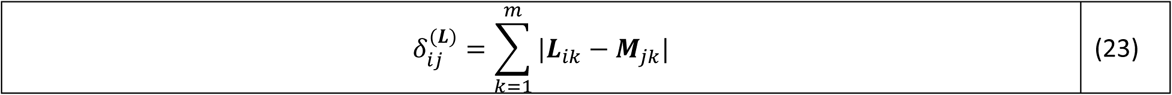

where *m* represents the number of variants and ***L***_*ik*_ represents the allele for *k*^*th*^ variant of *i*^*th*^ individual in leakage panel. Similarly, ***M***_*jk*_ denotes the allele for *k^th^* variant of *j*^*th*^ metasample. For each metasample the average allele distance is computed by averaging the distance among leakage samples:

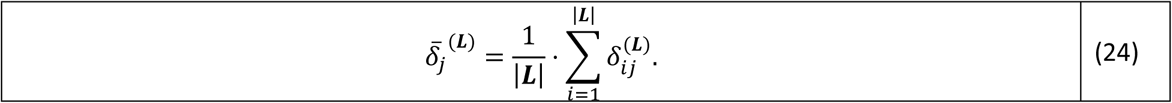

̅The average allele distance is computed between the metacohort variant alleles and the leakage panel and also the metacohort panel itself, i.e.

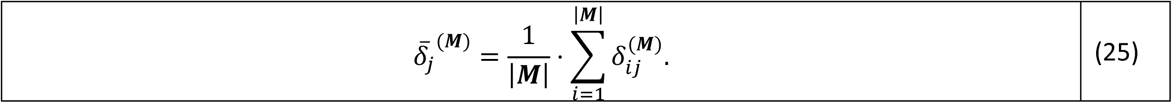

In addition, the minimum of the allele distances are used as statistics that measure the minimum distance between the metacohort sample variant alleles and the leakage estimation panel alleles:

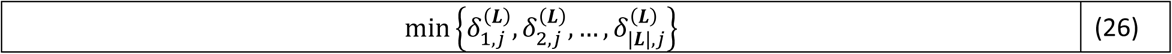

And the minimum distance between the metacohort sample alleles and the metacohort sample alleles:

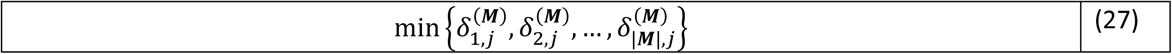

In the leakage statistics distribution plots (e.g. Fig 3b), we denote these statistics with “Min. Allele Distance”, “Avg. Allele Distance”, “Min. Meta Allele Distance”, “Avg. Meta Allele Distance”.

##### Allele Match Run Length Statistics

The number of consecutive alleles with exact match between the metacohort and the leakage estimation panel is computed. For this, we compare the variant alleles of *j*^*th*^ metasample, i.e., ***M_j_***, and all the samples in ***L*** and identify the longest-match run among alleles:

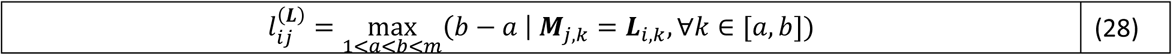

where 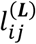 denotes the maximum match length between *j*^*th*^ metasample and *i*^*th*^ leakage panel sample. We compute the maximum of the maximum match length over all the leakage panel samples to assign the maximum match length for *j*^*th*^ metasample.

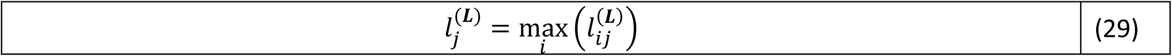

We defin 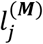 similarly by comparison of the metasamples with each other. These statistics are denoted by “Longest Match Length” and “Longest Meta Match Length” in leakage statistic distribution plots (e.g. Fig 3b).

#### Allele Frequency Based Statistics

The allele frequency-based statistics aim to quantify whether the hidden individual is at the extremes of allele frequency of their alleles within the metacohort. For this, we compute, for each metasample, the several statistics related to the allele frequency of the variants. These statistics rely on the allele frequency of the variants:

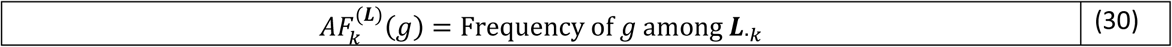

Where 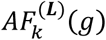 denotes the allele frequency of *g* for the *k*^*th*^ variant within the leakage panel. ***L***_·k_ denotes the vector of alleles at k^*th*^ variant in leakage panel. For each metasample, we compute the average, mean, and standard deviation of the allele frequency estimated from leakage panel and the metacohort variant alleles:

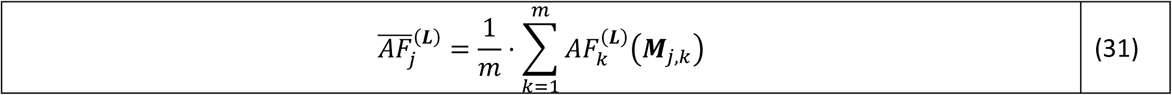

The standard deviation is defined as:

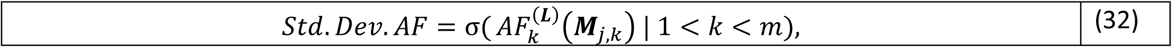

where *σ* denotes the standard deviation of the allele frequencies of the *m* variants. The minimum allele frequency statistic is computed as:

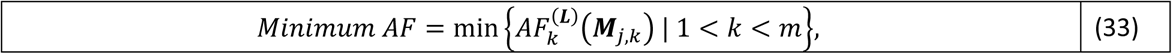

Similarly, the maximum allele frequency statistic is computed as:

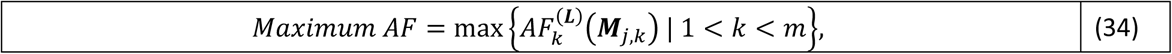

The allele frequencies statistics presented above are computed with respect to leakage panel. We also compute the allele frequency statistics with respect to the frequencies derived from the metacohort samples, i.e., 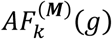. These are computed by replacing 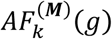 in the above formulas.

These statistics are denoted by “Min. AF”, “Max. AF”, “Avg. Meta AF”, “S.D. Meta AF”, “Avg. AF”, and “S.D. AF” in leakage statistic distribution plots.

#### Markov Chain Probability Based Statistics

For each metasample, we compute the aggregate Markov chain probability as

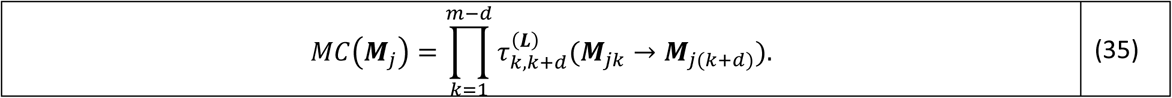

*MC*(***M***_*j*_) denotes the aggregate Markov Chain probability for *j*^*th*^ metasample. *d* represents distance between consecutive variants. This equation computes the aggregate transition probability between consecutive variants that are *d* indices apart from each other. By tuning *d*, the higher order correlation between variant alleles are incorporated in the statistic. 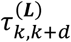 denotes the matrix of transition probabilities from allele ***M**_j,k_* of *k*^*th*^ variant to allele ***M***_*j*,(*k*+*d*)_ of (*k* + *d*)^*th*^ variant. This matrix is estimated from the leakage estimation panel ***L*** for transitions between consecutive variants. To evaluate the higher order statistics, we use higher order dependencies by changing the distance between the variants, *d*. In the leakage statistics The Markov Chain based statistics estimate the higher order composition of the variant alleles in each metasample. This enables measuring the leakage statistics that stem from the dependence of the consecutive variants.

The transition probabilities of the Markov Chain are estimated using the frequency of transitions observed in ***L***.

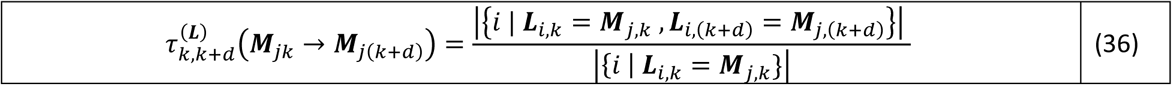

The transition probabilities are estimated from as the ratio of number of transitions divided by the frequency of observing ***L***_*i,k*_ = ***M***_*j,k*_ in ***L***. We use *d* = {1,2,3,4,5,10} as representative values of distance between variants while computing Markov chain probabilities.

##### Number of Zero Transitions in Metacohort Samples

This statistic is computed for each metasample by counting the number of transitions with exactly 0 transition probability:

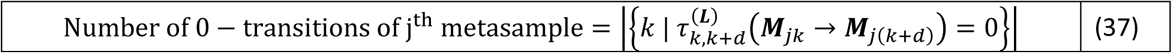

This statistic quantifies the number of transitions between consecutive alleles that have are never observed in the leakage quantification sample. This is the only statistic that evaluates the existence of de-novo variants within the metacohort.

These statistics are denoted by “1^st^ MC Prob.”, “2^nd^ MC Prob.”, “3^rd^ MC Prob.”, “4^th^ MC Prob.”, “5^th^ MC Prob”, “10^th^ MC Prob”, and “Zero Transitions” in the leakage statistics distribution plots. Each “MC Prob.” statistic corresponds to the order, i.e., *d* in (35), at which the dependency between the variants.

### Rank Normalization of the Leakage Statistics Matrix

In the tests, we computed the leakage statistics for each metasample in each hiding (Fig 2a). This yields a matrix of where the rows are the samples and the columns and the leakage statistics. Next, for each leakage statistic, we sorted the metasamples with increasing statistic. When there are ties, we assigned the average of the tied rank to all the samples. After the ranks are assigned, the leakage statistics matrix is converted into a statistic rank matrix for which each row is a sample and each column is a leakage statistic. Each entry contains the rank of the corresponding individual among all the individuals (rows) for the statistic. For each statistic, we recorded the rank of the hidden sample (i.e., the CEU samples in the tests)

### Joint Usage of Leakage Statistics via tSNE

We analyzed the rank normalized leakage statistics matrix using t-distributed stochastic neighbor embedding (tSNE)[48]. This analysis is performed for decreasing the dimensions of the leakage statistics. We used Rtsne package in R. We set the iteration number to 500 and perplexity to 50 while running tSNE.

## Acknowledgements

We thank Akdes Serin Harmanci for productive discussions about the conceptual development of the hiding framework.

## Supplementary Information for

We present supplementary text and figures.

### Importance of Generation Panel and How Well It Spans the Genotypic Space of Shared Variants

The most vital component in hiding based data sharing is how well generation panel represents the genotypic space of the variants that Alice wants to share. The generation panel serves as the knowledge-base that HaploHide uses to model the haplotype structure and allele frequencies of the variants. We know that the haplotype structure manifests the correlation between the alleles of variant in the vicinity of each other. For strong LD blocks, we employ the projection-based hiding so that the allelic structure is conserved by directly sampling the real individuals from the generation panel. For weak or no LD regions, there is no (or very low) variant-to-variant allele correlation. In this case we employ the sampling-based hiding and the variant alleles are generated independently from the allelic distributions learned from the generation panel.

One important observation is that the projection-based approach does not hide variants in weak LD blocks. While this is counter-intuitive, this result stems from the fact that projection creates a bias for variant selection. In particular, we observed that the projections tend to select variants that have high allele frequency while hiding variants in weak LD blocks. In this case, we can observe this variant selection bias in the aggregate Markov Chain probability-based leakage statistics. In other words, the generation panel cannot reliably represent the genotypic variation for variants that are in weak LD blocks. For strong LD blocks, the variant selection procedure does not create this bias because the generation panel sufficiently spans the possible set of variants on the region. Thus, the projection does not create a bias, i.e., it can capture the variant alleles of a new individual sufficiently well. We demonstrated this while hiding the variants in genes. These highlight the vital importance of the generation panel and how diverse it is in capturing the genotypic variation in the set of variants that Alice wants to share.

An important question is this: How will Alice share a set of variants such that some of them are in strong LD blocks and some of them are in weak LD blocks? In this case, there can be two ways of hiding. First is dividing the variants with respect to whether they are in high and low LD blocks and sharing the variants in different metacohorts such that high LD variants are hidden using projection-based approach and low LD variants are shared using sampling-based approach. We foresee that more complex models for haplotype generation can be employed (such as [[cite]]) to hide the variants with mixed LD characteristics and these will enable generating an effective crowd. We are not considering these approaches in the current study for the sake of simplicity of the presenting the hiding-based genomic data sharing.

#### The Probability Distribution of Metacohort Samples

We semi-quantitatively analyze the probability that an individual in the generation panel is observed in the metacohort. Given the ancestral panel of the individuals, and the generation panel, the probability that a specific individual is in the metacohort is:

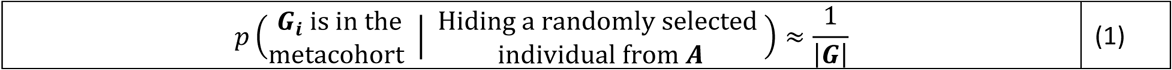

Where ***G_i_*** denotes *i^th^* individual in the generation panel. As in the main text, 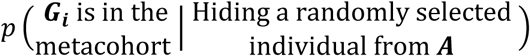 denotes the probability that ***G_i_*** is output in the metacohort while a randomly selected individual from ***A*** is being hidden.

We consider two cases depending on whether ***G_i_*** is a metasample or it is the hidden individual:

##### G_i_ is a metasamples in the metacohort

In the projection-based hiding, all the individuals other than the hidden individual (i.e., metasamples) are randomly sampled from the generation panel, ***G*** with exactly 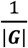 probability of being selected. Thus, if ***G_i_*** is a metasample, the probability of observing it in the metacohort is exactly 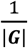.

##### G_i_ is the hidden individual in the metacohort

If ***G_i_*** is the hidden individual, this means that a randomly selected individual from ***A*** was projected onto ***G_i_***. In this case, we consider the probability of selection from ***A*** and selection of samples among the projected samples separately. For this, we first define the subset of individuals in ***G*** whose haplotypes are identical to the haplotype of ***G_i_*** and denote the set of these individuals with ***ã***. Next, we define the set of individuals in A, whose projections onto ***G*** is ***ã*** and we denote this set with ***a***. It should be noted that individuals in ***a*** have very similar haplotypes since their projections are exactly the haplotype shared by individuals in ***ã***. (Supplementary Fig. 2a)

In other words, projection of any individual in a is a randomly selected individual in ***ã*** and ***G_i_*** is an individual in ***ã***. Thus, a represents a set of individuals whose projections on ***G*** is any one of the individuals in ***ã***, which include ***G_i_***. Therefore, the hidden individual must be in the set a. Moreover, after projection, ***G_i_*** must be selected as the individuals among |***ã***| individuals. Putting these together:

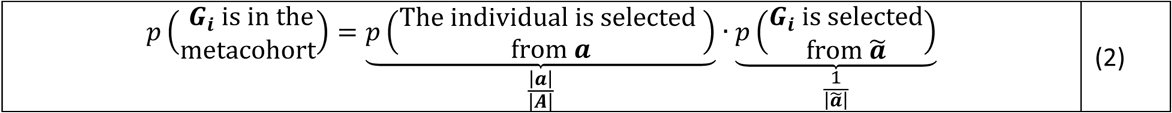

or simply as:

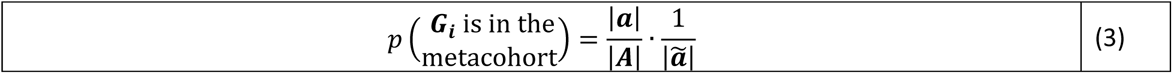

The first fraction in (3) represents the probability that an individual in ***a*** alleles is selected from ***A***. The second fraction in (3) represents the probability that we select ***Q_i_*** from among the individuals in the set ***ã***(Supplementary Fig. 2a).

Combining (3) with (1), the hiding condition in (1) can be summarized as:

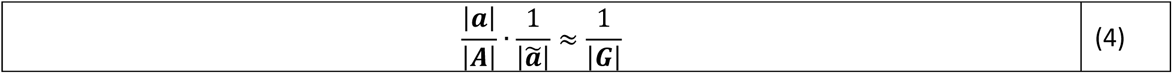

We can reorder this as:

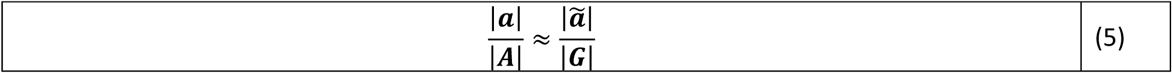

This condition implies that the proportion of individuals in ***G*** whose haplotype alleles match ***G_i_*** should be the same as the proportion of individuals in ***A*** whose haplotypes get projected on ***ã***. Alternatively, (5) refers to the fact that the haplotype frequencies of A and ***G*** match each other. Since we are interested in a general condition for everyone (not just ***G_i_***), this condition must hold for every individual (i.e., every haplotype) in ***G***. Consequently, the uniform probability distribution of metacohort individuals requires that the haplotype frequency distributions in ***A*** and ***G*** must match. This holds if the genetic background of the individual to be hidden matches or is very close to the genetic background represented by ***G***.

**Supplementary Figure 1:**
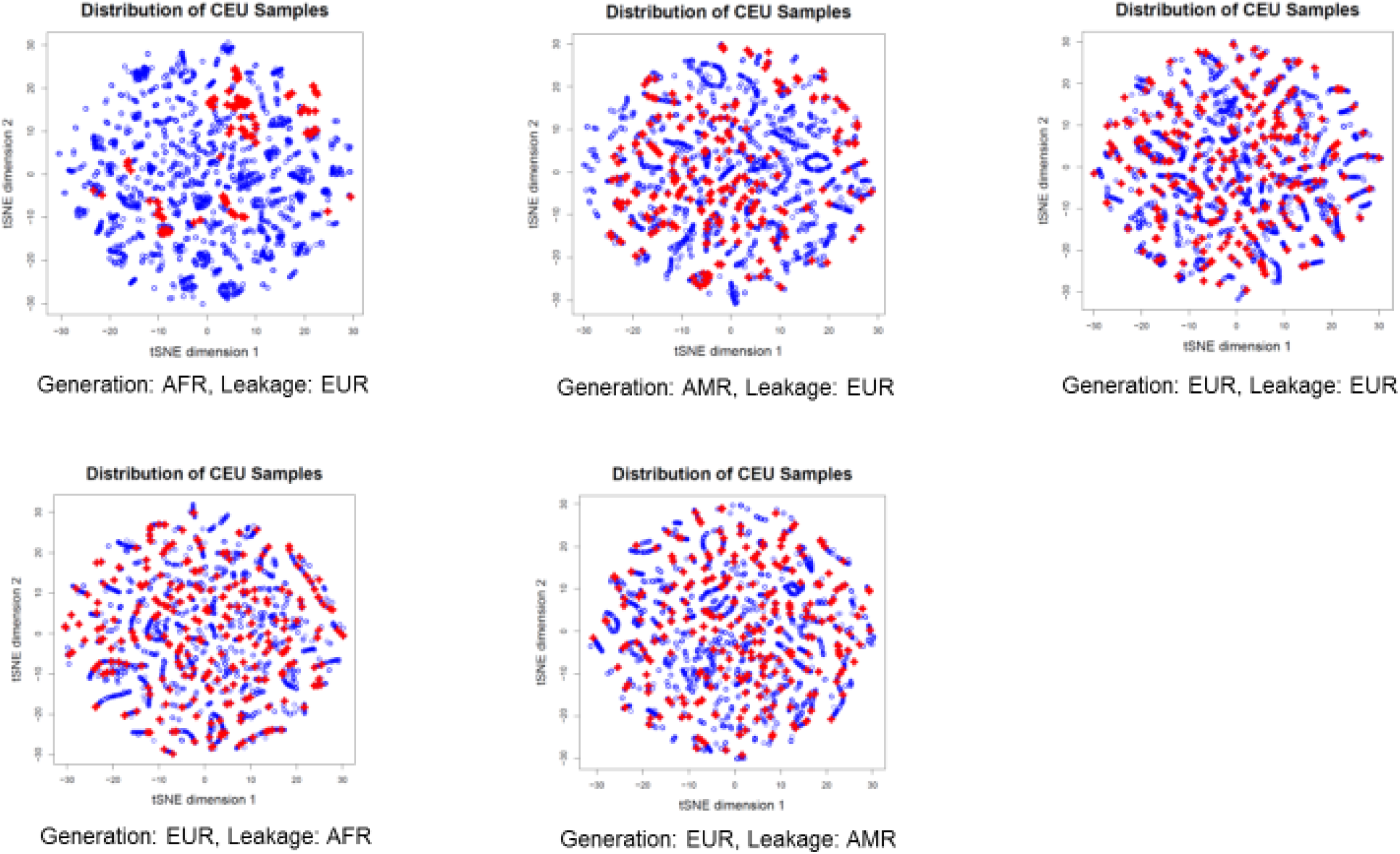
Scatter plot of t-statistic neighbor embedding of hidden individuals and other metasamples using the leakage statistic values. Each dot represents a sample such that red dots represent the hidden samples and blue dots represent other meta samples. Each scatter plot corresponds to different selection of populations for generation and leakage estimation panels indicated below the plots.

**Supplementary Figure 2:**
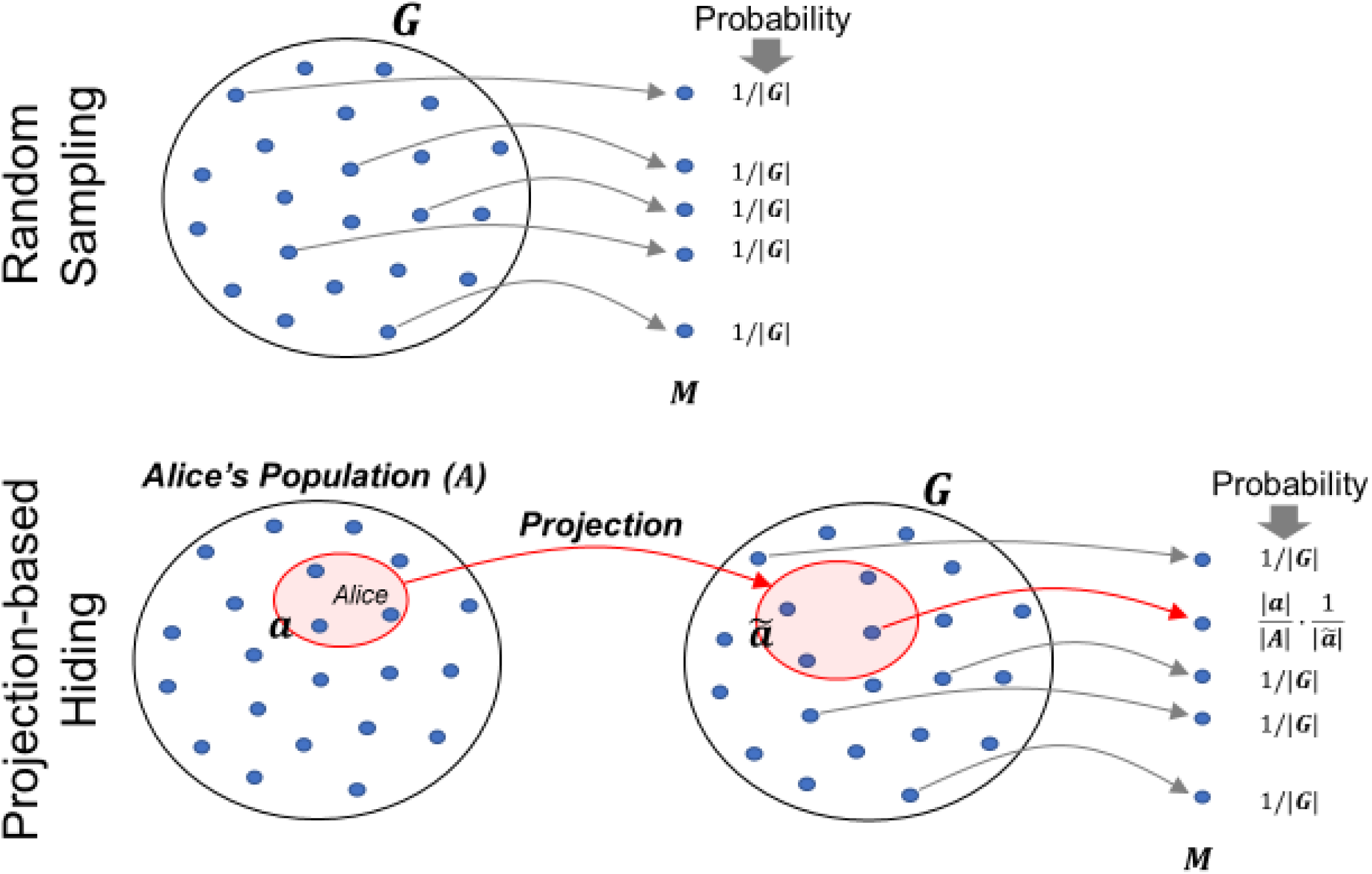
Illustration of the comparison of probability of generating 5 samples by random sampling from the generation panel ***G*** (above) and by projection-based hiding (below). On the right side, probability of generating each sample is shown. For random sampling, each sample is generated with *1/|**G**|* probability where *|**G**|*denotes the number of individuals in in the generation panel ***G***. For projection-based hiding, all the individuals other than the hidden individual are sampled randomly from ***G***. Thus, for these individuals are sampled with probability *1/|**G**|*. The hidden individual originates from the ancestral population ***A***. Thus, the hidden individual is first randomly sampled from the ancestral population ***A***. Then the selected individual is projected onto ***G*** and individual is selected among the projected set. Here, ***a*** denotes the set of individuals in ***A*** whose projections are denoted by ***ã*** in ***G*** such that the hidden individual’s projection is shown in the figure. Thus, the probability of selecting this individual is 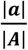. 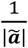 as shown in the Equation (5).

